# HCM-mutation G716R in β-myosin alters contractile function and responsiveness to mavacamten of human ventricular myofibrils

**DOI:** 10.64898/2026.11.19.767856

**Authors:** Anja Brunkhorst, Tim Holler, Birgit Piep, Antonio Francino, Cristobal G. dos Remedios, Theresia Kraft, Bogdan Iorga

## Abstract

Effects of the hypertrophic cardiomyopathy (HCM) associated mutation Gly716Arg in β-myosin at the sarcomeric level are unresolved. To further understand HCM-pathomechanisms in relation to modern treatment, we investigated the effect of this mutation in subcellular myofibrils isolated from HCM-patient’s myocardium compared to myofibrils from healthy donor hearts, and tested how mavacamten influences force-related parameters. Mavacamten is a state-of-the-art myosin inhibitor aimed to treat ventricular hypercontractility of symptomatic HCM patients.

Compared with donor myofibrils, patient myofibrils developed similar isometric forces, while kinetics of maximum force generation and particularly of relaxation were slower. Relaxation impairment was exacerbated when patient myofibrils were incompletely relaxed to mimic residual diastolic calcium levels. Inhibitory effects of mavacamten were dependent on force activation level and significantly less efficient in patient myofibrils. While the drug accelerated full relaxation of donor myofibrils, the slowing down effect of the mutation was not reversed by the drug in patient myofibrils. At partial relaxation, impairment was further aggravated by mavacamten in patient myofibrils.

This study suggests that HCM-associated mutation G716R in β-myosin does not evoke gain-of-function in human left ventricular myofibrils, and impaired relaxation could not be rescued by mavacamten. Our findings suggest that HCM-associated mutations may modulate myofibrillar sensitivity to mavacamten, providing a potential mechanistic explanation for differences in treatment response.

Studies such as the present one highlight the importance of mutation-dependent sarcomeric properties in understanding variability in responses to myosin modulators and potentially informing more individualized treatment strategies.

## INTRODUCTION

Hypertrophic cardiomyopathy (HCM), a genetically heterogeneous cardiac disorder that follows an autosomal dominant inheritance pattern, is the leading cause of sudden cardiac death (SCD) in the young [26]. In the absence of other conditions, HCM is characterized predominantly by left ventricular (LV) and/or septal hypertrophy, often with diastolic dysfunction, variable fibrosis, cardiomyocyte and myofibrillar disarray, and, in many cases, LV outflow tract (LVOT) obstruction. Clinical severity ranges from mild symptoms to pharmacotherapy-refractory heart failure requiring transplantation [26]. Approximately 50% of genotyped HCM patients were found to carry a pathogenic DNA variant. Usually these are heterozygous mutations in a sarcomeric gene, most prevalent in *MYBPC3* and *MYH7* [16, 26]. Among *MYH7* mutations, the c.2232G>A (exon 19; p.Gly716Arg; ClinVar:14105) variant exhibits high penetrance and early-onset HCM across multiple ethnicities, with patients frequently developing asymmetric septal hypertrophy, arrhythmias, elevated SCD risk, and in some cases progressing to systolic dysfunction and heart failure requiring correcting interventions or transplantation [1, 17, 33, 46].

HCM currently is considered a disease of the sarcomere proposed to be driven by a hypercontractile mechanism, a concept that was experimentally corroborated by a series of studies examining missense mutations in βMyHC or other sarcomeric proteins [40]. One mechanistic hypothesis proposes that in HCM, mutated myosin heads are more likely to leave a putative sequestered, *off*-configuration of the myosin head (interacting heads motif, IHM) and become available to interact strongly with actin, thus causing an increase in force generation [13, 40]. Nevertheless, several HCM-linked mutations in βMyHC (such as converter domain mutations R712L-, R719W-, R723G-, and G741R-βMyHC) and other sarcomeric proteins produce rather a diminished calcium sensitivity and/or function-related kinetic parameters, contradicting the hypercontractile hypothesis [20, 24, 27, 39, 40, 45]. These findings underscore the variability in the functional consequences of HCM mutations, highlighting that ventricular hypercontractility is not universal and other mechanisms may co-exist.

The HCM missense mutation Gly716Arg (G716R) is located in the myosin converter domain, outside the putative IHM (Fig. 1A,B). The converter couples the motor catalytic core to the lever arm that rotates ∼70° during the power-stroke, being essential for force generation [14, 20, 34]. The G716R substitution, previously investigated in myosin of *Dictyostelium discoideum* (A699R) using the laser trap system, *in vitro* motility assay and an ATPase activity assay, weakened the binding affinity of the mutant myosin to actin and reduced the myosin-actin force [12]. Yet, despite genetic and clinical characterisation of some patients with this mutation, the primary functional effect of the G716R-βMyHC mutation on sarcomeric performance in human myocardium remains unresolved.

**Figure 1.**
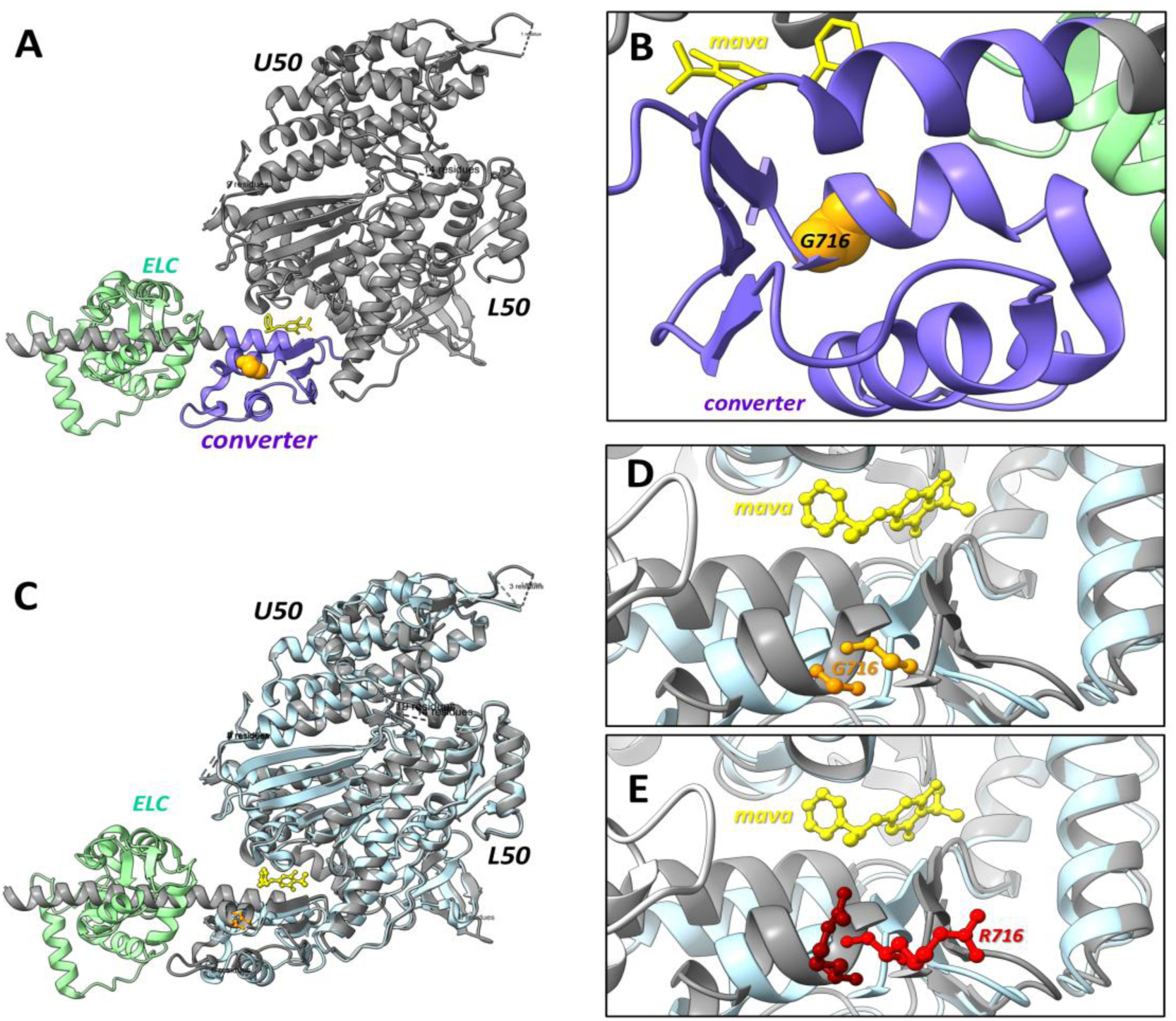
Structural elements related to G716R substitution and mavacamten in myosin. **A**. Elements of the secondary structure (grey and light-green for the *ELC*) of the β-myosin S1-fragment in the pre-power-stroke state (PPS-S1-Mava, chain-A: 3-807 residues, PDB-ID <u>8QYQ)</u> [3] complexed to mavacamten (*mava*, yellow) and Mg·ADP·BeFx (not shown). Mava binds S1 at the interface between the converter (blue) and upper-50kDa (*U50*) domains. G716 (orange) is located in the converter, at a larger distance from mava than distances between mava and its interacting residues (L120, Y164, D168, H668, N711, R712, R721) [3, 29]. **B**. Detail rotated with 180° of the converter with G716 and mava. **C**. Elements of the secondary structure (<u>8QYQ;</u> grey) of myosin-S1 complexed to mava superimposed with the structure in absence of mava (PPS-MD-Apo structure, chain-A: 32-780 residues, light-blue, PDB-ID <u>8QYP)</u> [3] in the pre-powerstroke state complexed to Mg·ADP·Vanadate (not shown). **D**. Detailed view (*ELC* in the left side is not shown) indicating mava-induced structural shift in the backbone and position of the G716 (orange in 8QYQ +mava, dark-orange in 8QYP -mava). **E**. A similar detailed view as in D, in which G716 was replaced by R716 (red in 8QYQ +mava, dark-red in 8QYP -mava); nevertheless, the indicated structural shift represents only mavacamten binding, not the potential allosteric shift caused by the 716G>R mutation itself. Abbreviations: *ELC*, essential light chain; *U50*, upper-50kDa domain of myosin-S1; *L50*, lower-50kDa domain of myosin-S1; *converter*, converter domain of the myosin-S1. Images were generated with <u>ChimeraX.</u>

Meanwhile, treatment with the myosin motor inhibitor of a specific group of HCM patients with obstructive phenotype is quite well established [28, 32]. *In vitro* studies in presence of mavacamten, mostly with WT myosin-based preparations, indicated reversible reductions in ATPase activity, isometric force, and their Ca^2+^-sensitivity, but also an accelerated relaxation and a preserved length-dependent activation [21, 31, 36]. These findings were explained with mavacamten mainly inhibiting phosphate (P_i_) release during ATP turnover and stabilizing the myosin *off*-state, thereby limiting the pool of heads strongly-bound to actin, participating in force generation [2, 21, 31, 35].

β-cardiac myosin heads reversibly bind mavacamten between the N-terminal region of its upper-50kDa subdomain and the converter region (Fig. 1A,B) [3, 8]. Since it cannot be ruled out that some myosin mutations may impede the proposed action of mavacamten, it is of interest to understand the effect of this drug on sarcomeric function in presence of individual HCM mutations, as well.

In this study we used human ventricular myofibrils (hvMFs) isolated from a heterozygous HCM patient’s heart carrying the G716R-mutant myosin (MT-hvMFs) and hvMFs isolated from donor hearts (WT-hvMFs), considered as reference, to investigate the primary functional effect of the G716R-βMyHC mutation on sarcomeric performance and its sensitivity to mavacamten treatment. The latter aims to understand whether the drug normalizes sarcomeric function under coexistence of both, MT-and WT-myosin heads in MT-hvMFs of the heterozygous patient. Main findings were that the HCM-linked G716R mutation does not cause sarcomeric hyperfunction but impairs cross-bridge cycling and myofibrillar relaxation. Surprisingly, mavacamten did not rescue the impairment.

## MATERIALS and METHODS

### Experiments design

Three categories of micromechanical experiments were performed on human ventricular myofibrils: (**a**) The first approach basically characterized patient-derived myofibrils relative to donor-heart references (Fig. 4,5; Table S1A), complemented by isoform analyses of major sarcomeric proteins involved in force generation, regulation, and modulation (Fig. 3), and by histology and immunostaining of thin ventricular tissue slices (Fig. 2); (**b**) The second approach focused on force-related kinetic parameters during myofibrils relaxation in the presence of residual [Ca²⁺] (Fig. 6; Table S1B); (**c**) The third one investigated the effects of mavacamten on myofibrillar contraction-relaxation at maximal Ca²⁺-activation followed by either full or partial relaxation (Fig. 7,8).

**Figure 2.**
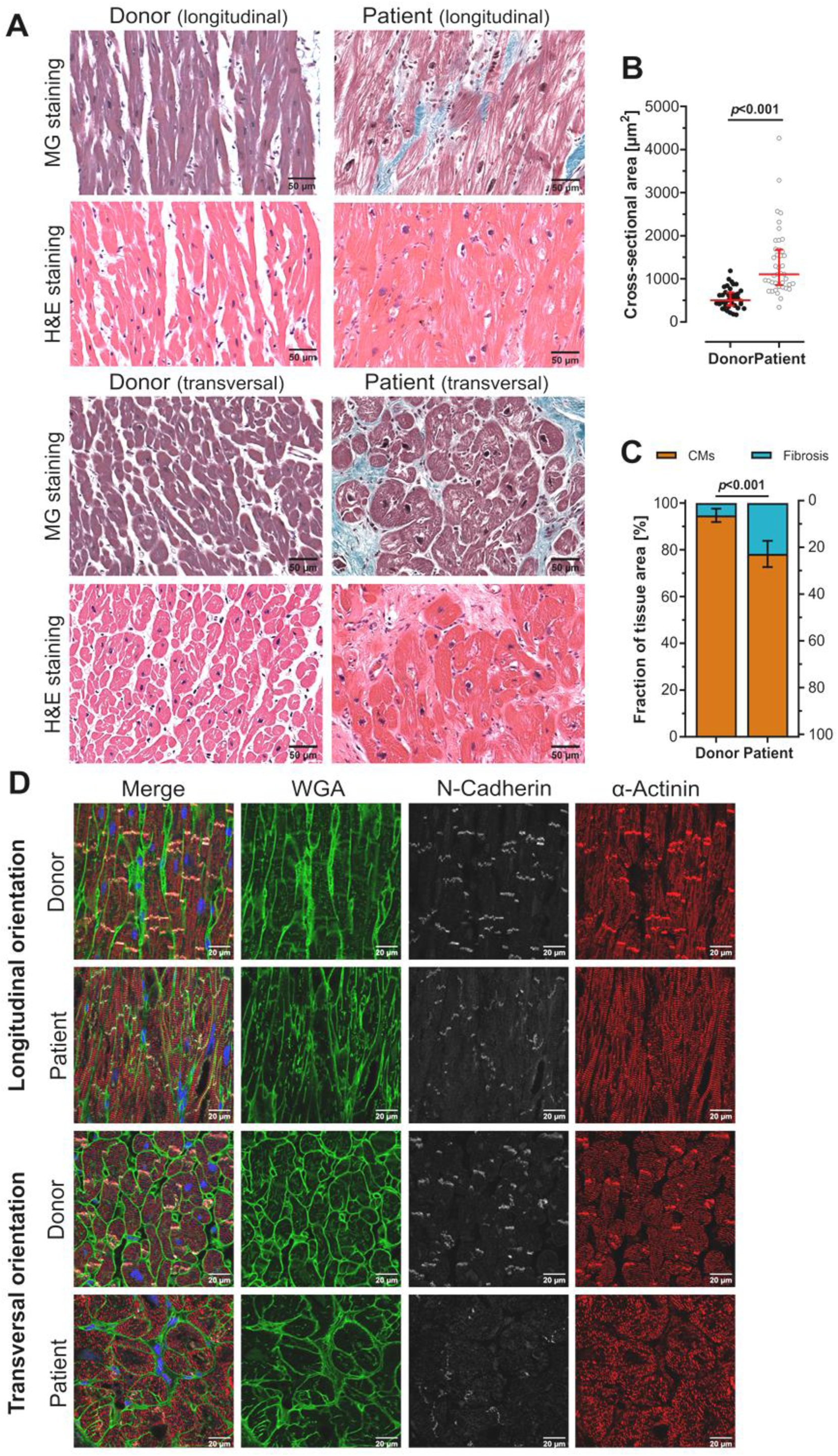
Histological and immunofluorescence analysis of patient and donor left ventricular tissue. **A**. Representative histological sections of donor and patient left ventricular (LV) wall stained with Masson-Goldner (MG) and haematoxylin and eosin (H&E); *top*: overview of tissue morphology in longitudinal section showing cardiomyocytes (CMs) area; *bottom*: quasi-transversal section from which cross-sectional-area (CSA) of CMs was estimated. **B**. For CSA analysis, a total tissue area of 2.63 mm² was evaluated in donor and 11.44 mm² in patient samples; analyses were based on 8 donor and 3 patient tissue LV sections; scatter dot plot depicts individual cell measurements analyzed from different regions of 2.63 mm^2^ donor and 11.44 mm^2^ patient LV tissue, with median and interquartile range indicated; statistical significance was assessed using the Mann-Whitney *U*-test (*p*<0.0001). **C**. Fibrosis quantification from MG-stained sections was analyzed from 7 different donor and 3 different patient LV tissue sections; for each donor or patient, a total tissue area of 166,944 µm² (n=8 regions of interest) was analyzed. Stacked bar plot shows fractional area of fibrosis-stained regions and cardiomyocytes, with mean ± standard deviation (SD); statistical significance was assessed using an unpaired Student’s *t*-test (*p*<0.0001). **D**. Representative immunofluorescence images of donor and patient myocardium: left-to-right, merge image with localization of all markers and DAPI (nuclei; blue), WGA (cell borders; green), N-Cadherin (intercalated discs; white), and α-Actinin (sarcomeric pattern; red).

**Figure 3.**
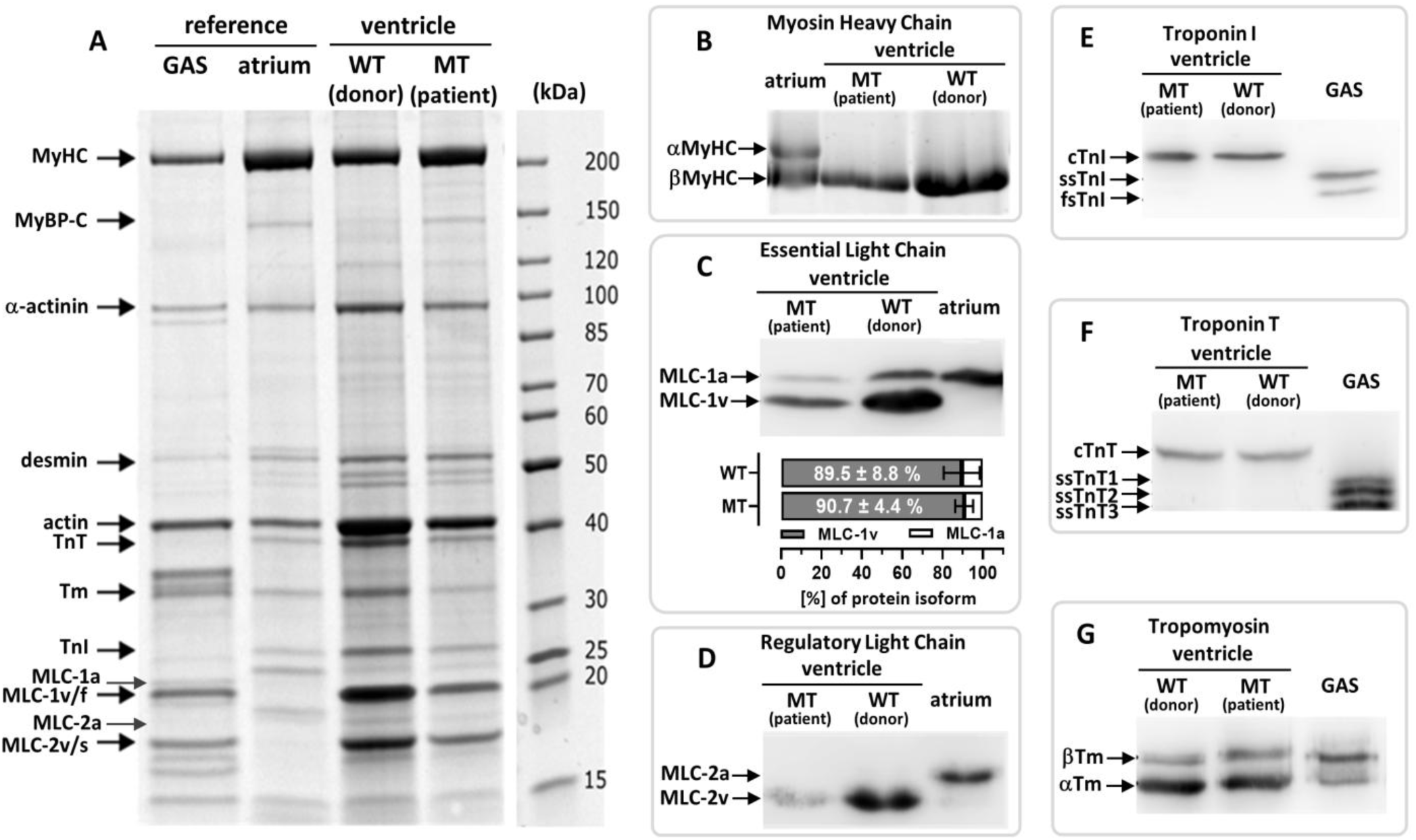
Sarcomeric protein isoform profile of patient (MT) and donor (WT) left ventricular myofibrils. Coomassie-stained SDS-gel identifies several proteins and their isoforms: **A**. α-actinin, desmin, actin, troponin T (TnT), tropomyosin (Tm), troponin I (TnI), myosin binding protein C (MyBP-C), essential light chain (MLC-1), regulatory light chain (MLC-2); **B**. Myosin heavy chain (MyHC) while βMyHC isoform predominated in ventricular samples. Western Blot analysis was used to identify: **C**. Atrial (MLC-1a) and ventricular (MLC-1v) isoforms of MLC-1; **D**. Atrial (MLC-2a) and ventricular (MLC-2v) isoforms of MLC-2; **E**. Cardiac (cTnI), slow (ssTnI) and fast (fsTnI) skeletal muscle isoforms of TnI; **F**. Cardiac (cTnT) and three distinct slow (ssTnT_1_, ssTnT_2_, ssTnT_3_) skeletal muscle isoforms; **G**. Alpha (αTm) and beta (βTm) isoforms of Tm. Myofibrils isolated from human atrium and skeletal muscle gastrocnemius (GAS) tissues served as references; in the GAS lane, the bands of MLC-2s and MLC-1f (or MLC-1sb) skeletal muscle isoforms are indicated. Horizontal bar plot (**C bottom**): proportions of the atrial (open) and ventricular (grey filled) MLC-1 isoforms; mean±SD, n=5 (WT), n=4 (MT); no significant differences (*p*>0.05; Student’s *t*-test); myofibrillar samples taken from patient and three pooled donor tissues.

**Figure 4.**
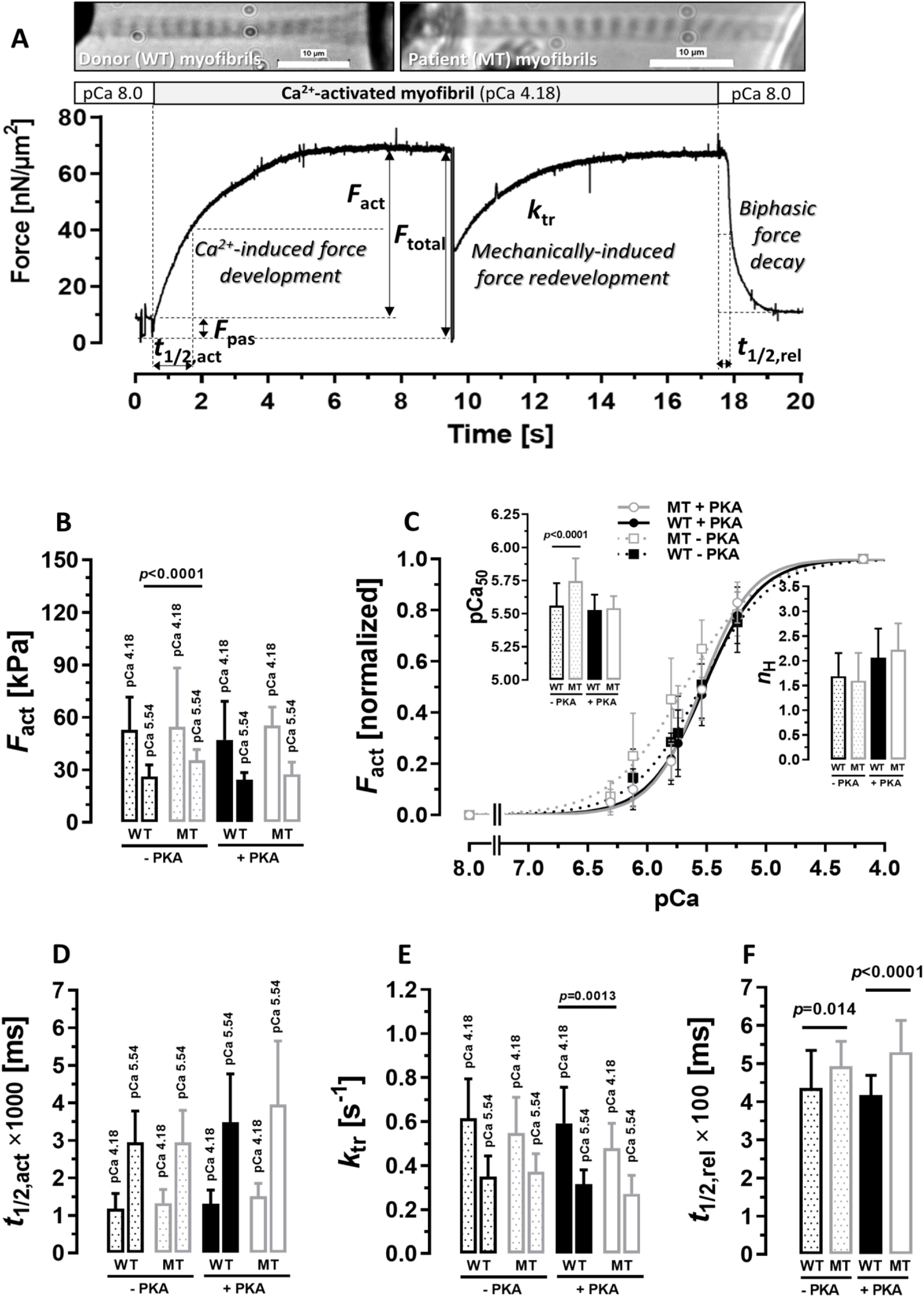
Force-related parameters of patient and donor myofibrils. **A**. Top: Images for donor WT-hvMFs (**left**) and patient MT-hvMFs (**right**). Bottom: Representative force trace of contracting-relaxing WT-hvMFs indicating force-related parameters detailed in *Supplementary Materials*: passive force (*F*_pas_), activation half-time (*t*_1/2,act_) of Ca^2+^-induced force development, rate constant (*k*_tr_) of force redevelopment, total isometric force (*F*_total_) and the relaxation half-time (*t*_1/2,rel_). Force-related parameters of WT-hvMFs (black dots-patterned or black filled bars) and MT-hvMFs (grey dots-patterned or grey open bars), before (dots-patterned bars, -PKA) and after (filled or open bars, +PKA) incubation with PKA: **B**. Maximal and submaximal force *F*_act_ (*F*_act_=*F*_total_-*F*_pas_; 1kPa=1nN/µm^2^); **C**. Normalized (=*F*_act_/*F*_act,max_) force-pCa relationship for WT-hvMFs (black filled symbols and curves) and MT-hvMFs (grey open symbols and curves), treated (+PKA, circles and solid curves) or not (-PKA, squares and dotted curves) with PKA, providing calcium sensitivity of force (pCa_50_; left inset) and the steepness (*n*_H_; right inset). **D**. *t*_1/2,act_ at pCa 4.18 and pCa 5.54. **E**. *k*_tr_ at pCa 4.18 and pCa 5.54. **F**. *t*_1/2,rel_ of full relaxation. *F*_act_ (**B**), *t*_1/2,act_ (**D**), and *k*_tr_ (**E**) at pCa 4.18 and pCa 5.54 are significantly different (*p*<0.001; Student’s *t*-test; not indicated) for each myofibrils category (WT-hvMFs, MT-hvMFs, ±PKA). Significant difference in *F*_act_ at pCa 5.54 (**B**) and in pCa_50_ (**C**) between PKA-untreated MT-and WT-hvMFs. Error bars: standard deviation (SD). The number of myofibrils contributing to the each indicated parameter is given in Table S1A (*Supplementary Materials*). WT-hvMFs prepared from three donor hearts; data are pooled. MT-hvMFs prepared from one HCM-patient heart.

**Figure 5.**
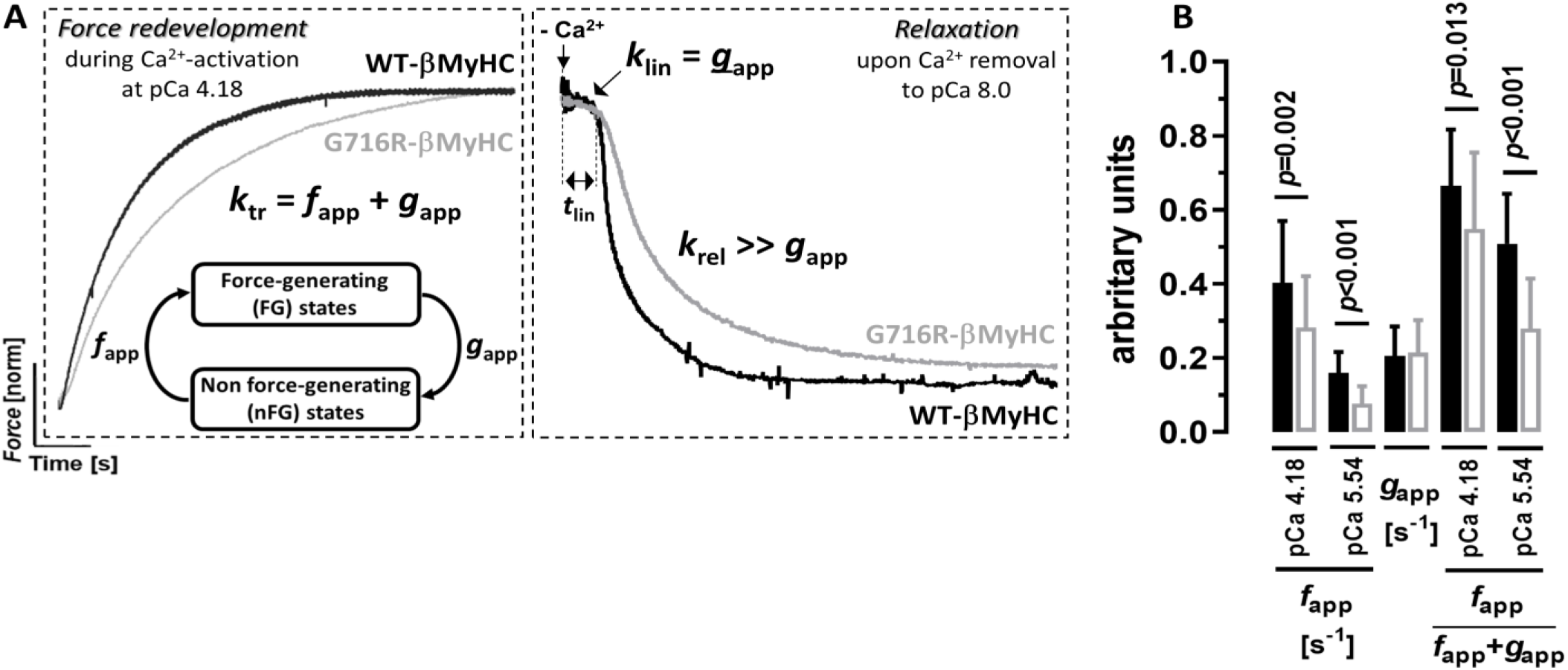
Cycling cross-bridge-related kinetic parameters of patient and donor myofibrils. **A**. Representative force traces reflecting kinetics (*k*_tr_) of force redevelopment of Ca^2+^-activated myofibrils at pCa 4.18 (**left**) and kinetics (*k*_lin_, *t*_lin_, *k*_rel_) of biphasic force decay upon sudden Ca^2+^ removal from pCa 4.18 to pCa 8.0 (**right**) for PKA-treated patient MT-hvMFs (grey traces) compared with donor WT-hvMFs (black traces). *k*_tr_ depends on the rate constants *f*_app_ and *g*_app_ describing cycling cross-bridges that enter and leave force-generating states, respectively. 1^st^ relaxation phase is described by the rate constant *k*_lin_ and duration *t*_lin_; the 2^nd^ relaxation phase is described by the rate constant *k*_rel_. **B**. Bar plots indicate significant differences (*p*<0.05; Student’s *t*-test) of the *f*_app_ and duty-ratio *f*_app_/(*f*_app_+*g*_app_) between cross-bridge ensembles in MT-hvMFs (grey open bars) and in WT-hvMFs (black filled bars) at pCa 4.18 and pCa 5.54, respectively. There are no significant differences (*p*>0.05) for *g*_app_ between cross-bridges of MT-and WT-hvMFs. Error bars: SD. Number of myofibrils contributing to: *f*_app_ (n_WT_=30, n_MT_=35), *g*_app_ (n_WT_=65, n_MT_=35) and duty-ratio (n_WT_=30, n_MT_=35). Data from Table S1A.

**Figure 6.**
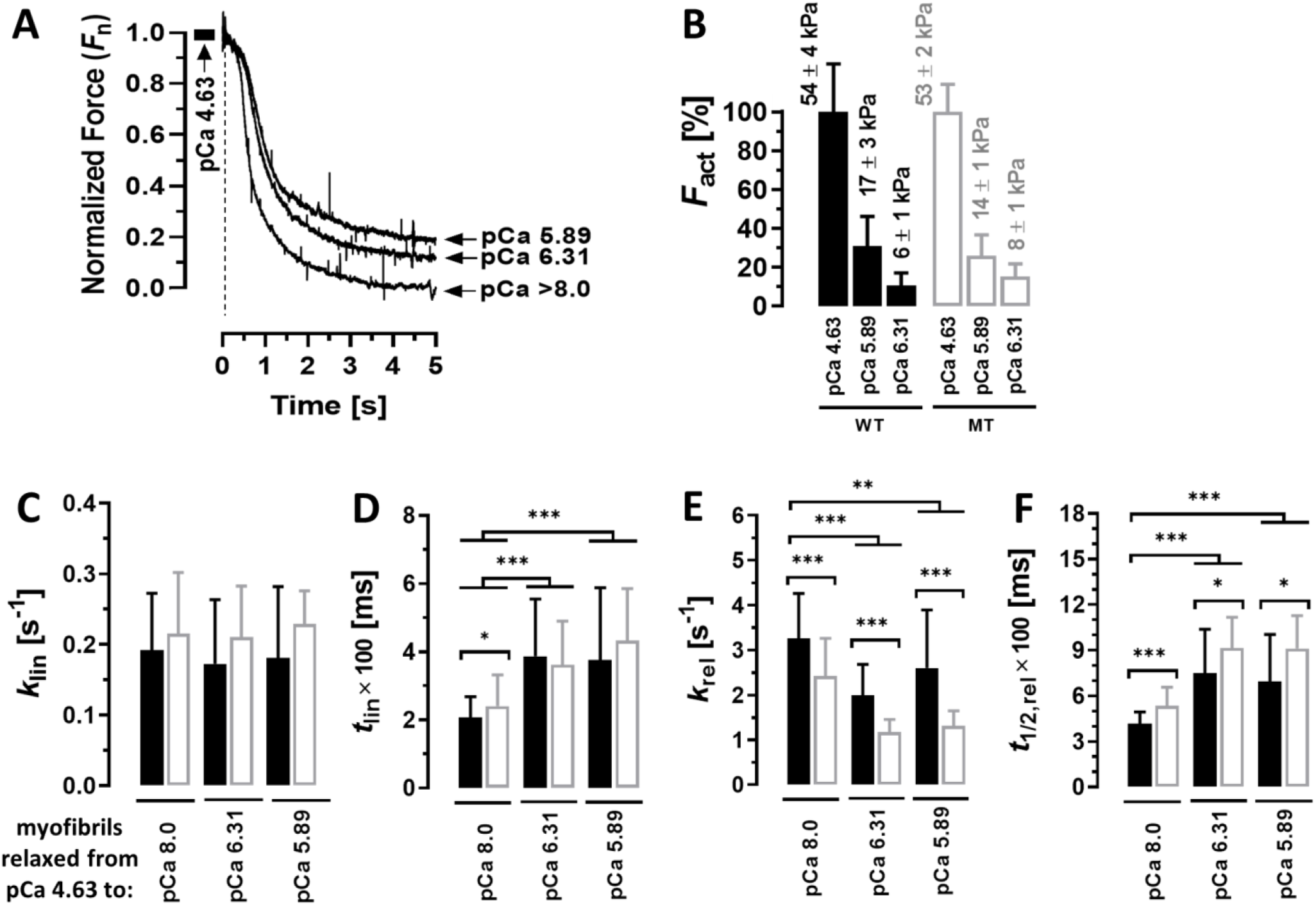
Kinetics of patient and donor myofibrils which relax at different calcium concentrations. **A**. Representative normalized force decay traces of myofibrils, which were initially Ca^2+^-activated at saturating [Ca^2+^] (pCa 4.63) and then either fully (at pCa 8.0) or partially (at pCa 6.31 or pCa 5.89) relaxed. The following parameters were compared between patient MT-hvMFs (grey open bars) and donor WT-hvMFs (black filled bars), and between partially relaxed and completely relaxed hvMFs: **B**. *F*_act_, isometric forces generated by hvMFs at partial Ca^2+^-activation levels (pCa 5.89, pCa 6.31) compared with *F*_act,max_ generated at saturating [Ca^2+^] (pCa 4.63). There were no significant (*p*>0.05) differences in force between MT-hvMFs and WT-hvMFs at each given [Ca^2+^]. *F*_act_ is given normalized to *F*_act,max_ in % and as mean±SEM. **C**. *k*_lin_, rate constant of the first relaxation phase. **D**. *t*_lin_, duration of the first relaxation phase. **E**. *k*_rel_, rate constant of the second relaxation phase. **F**. *t*_1/2,rel_, overall relaxation half-time. All myofibrils were incubated with PKA before functional analysis. Significance levels: *, *p*<0.05; **, *p*<0.01; ***, *p*<0.001; not significant, *p*>0.05 (Student’s *t*-test). Error bars: SD. Number of myofibrils contributing to each relaxation parameter at pCa 8.0 (for WT-hvMFs/MT-hvMFs): n_klin_=88/35, n_tlin_=110/37, n_krel_=110/37, n_t1/2rel_=88/35; The number corresponding to each parameter at pCa 6.31 and 5.89 is given in Table S1B (*Supplementary Materials*).

**Figure 7.**
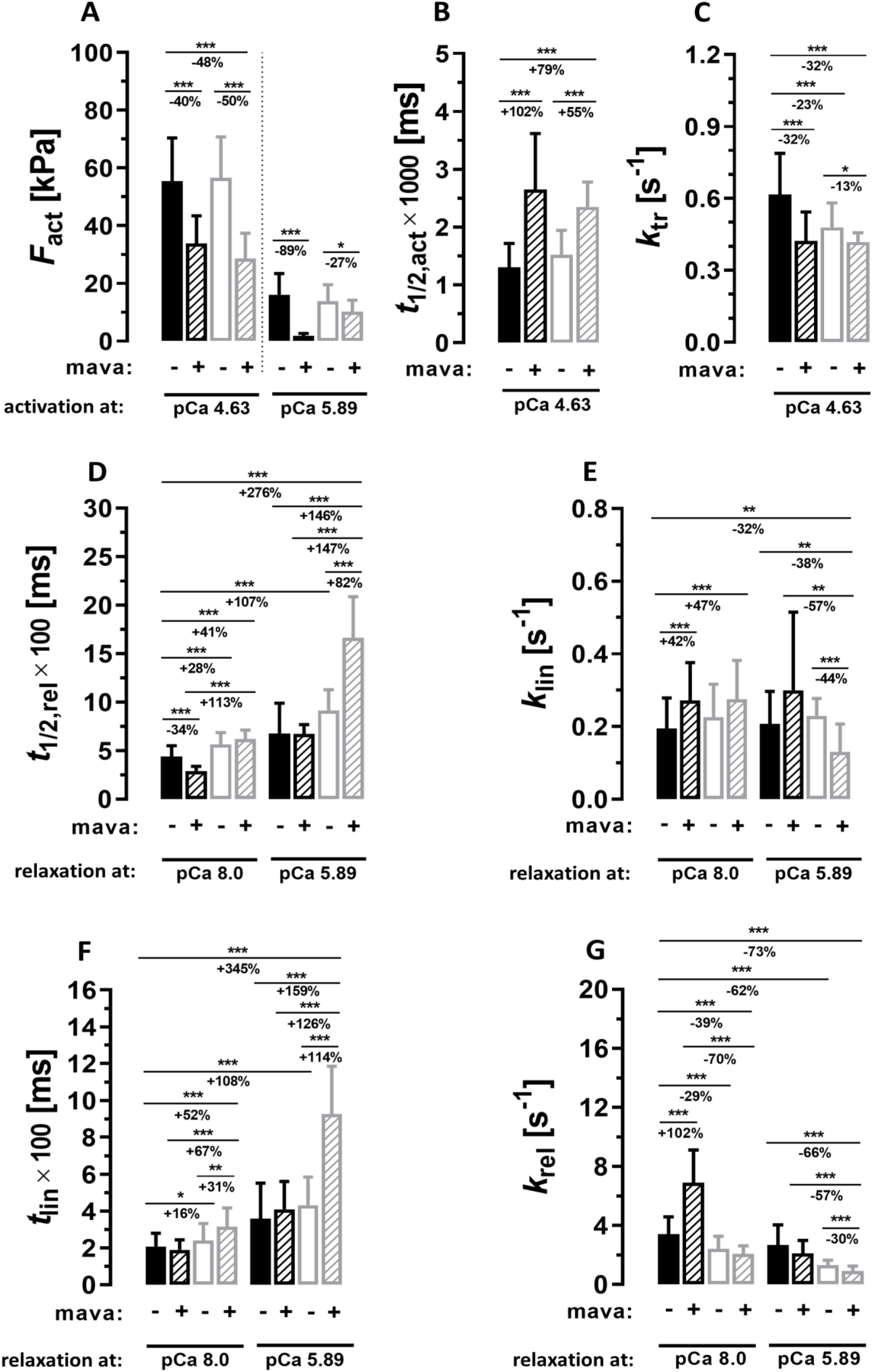
The effect of mavacamten on force-related parameters of patient and donor myofibrils. Mavacamten (mava) was present (+, patterned bars) or absent (-, not patterned bars) in activating (pCa 4.63 or pCa 5.89) and relaxing (pCa 8.0 or pCa 5.89) solutions to which PKA-treated MT-hvMFs (open grey or patterned grey bars) and WT-hvMFs (filled black or patterned black bars) were exposed. **A**. Generated isometric force (*F*_act_). **B**. Activation half-time (*t*_1/2,act_) of Ca^2+^-induced force development. **C**. Rate constant of force redevelopment (*k*_tr_). **D**. Relaxation half-time (*t*_1/2,rel_). **E**. Rate constant (*k*_lin_) of the first relaxation phase. **F**. Its duration (*t*_lin_). **G**. Rate constant (*k*_rel_) of the second relaxation phase. Percent (±%) indicates the parameter relative change when compared with the left side bar corresponding reference indicated by a horizontal line. Significance levels (Student’s *t*-test): *, *p*<0.05; **, *p*<0.01; ***, *p*<0.001; not indicated for *p*>0.05. Considering that a donor heart was not treated with mavacamten and its WT-hvMFs relax fully, comparison of the following data pairs in D, E, F, G were not shown: MT-hvMFs (-mava, relaxed at any pCa) with WT-hvMFs (+mava, relaxed at any pCa), MT-hvMFs (-/+mava, relaxed at pCa 8.0) with WT-hvMFs (-/+mava, relaxed at pCa 5.89), and WT-hvMFs (+mava, relaxed at pCa 5.89) with WT-hvMFs (-mava, relaxed at pCa 8.0). Error bars: SD. Number of investigated myofibrils Ca^2+^-activated at pCa 4.63 (or at pCa 5.89) and relaxed either fully at pCa 8.0 (n_WT-mava_=88 (40), n_WT+mava_=31 (8), n_MT-mava_=41 (19), n_MT+mava_=23 (20)) or partially at pCa 5.89 (n_WT-mava_=55, n_WT+mava_=7, n_MT-mava_=19, n_MT+mava_=20).

**Figure 8.**
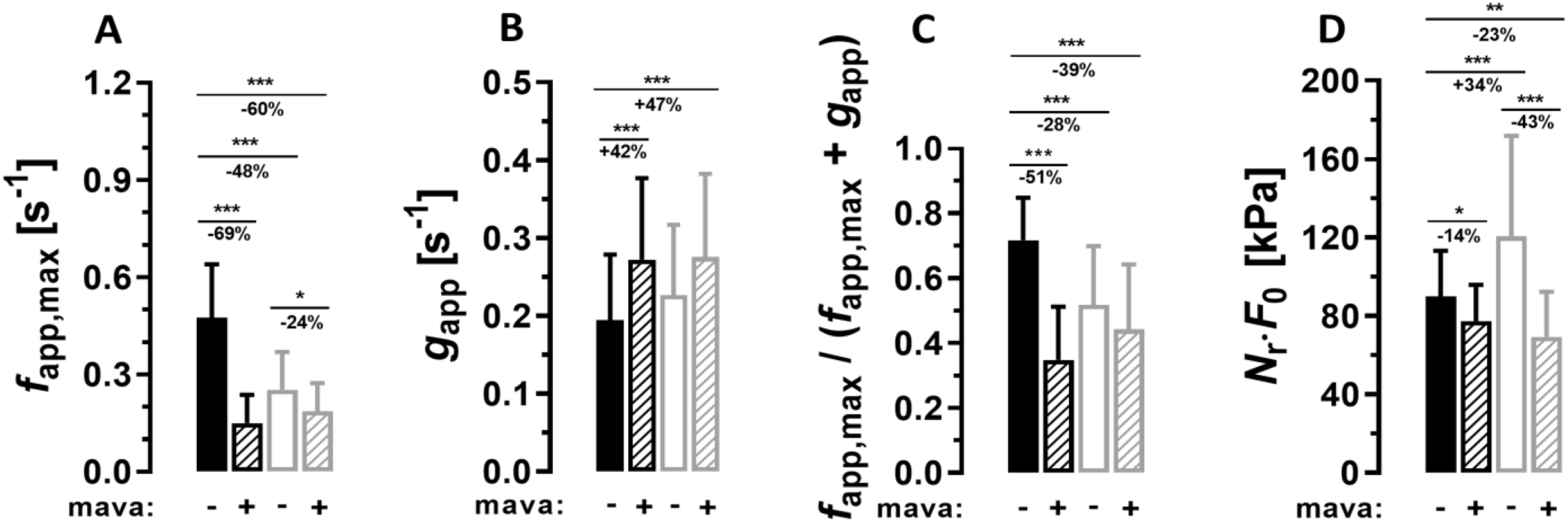
Effects of mavacamten on parameters related to the ensemble of cycling cross-bridges. Mavacamten (mava) was present (+, patterned bars) or absent (-, not patterned bars) in activating and relaxing solutions to which patient (open grey bars or patterned grey bars; MT) and donor (filled black bars or patterned black bars; WT) left ventricular sarcomeres were exposed. **A**. Maximum rate constant *f*_app,max_ of cross-bridges entering force-generating states in sarcomeres activated at pCa 4.63; **B**. Rate constant *g*_app_ of cross-bridges leaving force-generating states in relaxing sarcomeres at pCa 8.0; **C**. Maximum duty-ratio *f*_app,max_/(*f*_app,max_+*g*_app_); **D**. Cross-bridge *ensemble-force-factor N*_r_·*F*_0_. Meaning of percentage values and significance levels are detailed in the legend of Fig. 7. Error bars: SD. Number of myofibrils (WT-mava, WT+mava, MT-mava, MT+mava) contributing to each parameter, respectively: n_fapp,max_ = 55, 24, 39, 16; n_gapp_ = 88, 25, 39, 22; n_duty-ratio_ = 55, 24, 39, 16; n_NrF0_ = 55, 24, 39, 15.

### Patient data

Human myocardial samples were provided by Hospital Clinic, Cardiology Department, Barcelona (patient samples), St. Vincent’s Hospital in Sydney, Australia (donor) and Hannover Medical School (reference samples). Details about human samples and ethical statements are included in *Supplementary Materials*.

### Solutions Composition

Composition of solutions used to prepare (*Na-solution*, *K-solution*) and to activate (*Ca^2+^-activating-solution*) or relax (*Relaxing-solution*) myofibrils for investigations of their contractile function in presence or absence of mavacamten is described in *Supplementary Materials* and previously [19].

### Preparation of ventricular myofibrils

Details about myofibrils preparation from human myocardial (patient, donor) and skeletal muscular long-term cryopreserved tissues are described in *Supplementary Materials* and previously [19, 24].

### Samples treatments

Prior to investigations of contractile function, ventricular myofibrils were treated with protein kinase A (PKA) in most of experiments; if this was not the case, this is stated. In some experiments (e.g., Fig.7,8), mavacamten was present in both *Ca^2+^-activating-* and *Relaxing-solutions*. Details are given in *Supplementary Materials*.

### Histology and immunostaining

General tissue morphology, cross-sectional area (CSA) and fibrosis content assessments (Fig. 2A-C) were based on standard tissue staining; α-Actinin, N-Cadherin and WGA proteins (Fig. 2D) were identified based on immunofluorescent staining; details are described in *Supplementary Materials*.

### Analysis of sarcomeric proteins

Identification of some sarcomeric proteins mainly involved in force generation (myosin, actin), modulation (light chains) and Ca^2+^-regulation (troponin, tropomyosin) was performed using Coomassie-stained SDS-gels and Western Blots; details are in *Supplementary Materials*.

### Myofibrils micromechanical setup and derived parameters

Basic features and technical details of the custom-built micromechanical setup used to assess the contractile function of thin myofibrillar bundles are detailed in *Supplementary Materials* and previously [19].

### Data Analysis and Statistics

Description about data analysis, processing and statistics is detailed in *Supplementary Materials*.

## RESULTS

### Patient with moderate HOCM and functional impairments

The patient (male) carrying the myosin mutation G716R was diagnosed at age 20 with moderately obstructive familiar hypertrophic cardiomyopathy (HOCM, LVOT peak gradient 30-50 mmHg), exhibiting an early impairment of the infra-Hisian conduction and biventricular asymmetric hypertrophy (IVS thickness 15-19 mm), which was more severe for the left ventricle (LV). Ejection fraction was only slightly reduced (EF=40-45%). The initial functional capacity (NYHA class II) worsened progressively in the next 10 years. At the age of 27, persistent atrial fibrillation was observed accompanied by systolic dysfunction (EF=30-40%) and hypokinesia of the septum, apex and inferior segments.

At the age of 30, biventricular failure was caused by mixed severe systolic (EF=30%) and diastolic (restrictive, type-III pattern) dysfunction. Coronary heart disease, ischaemia and anemia were ruled out. Intraventricular obstruction decreased in parallel with progressive deterioration of myocardial systolic and diastolic function.

After the age of 30, the morpho-functional pattern resembled more to that found in patients with dilated cardiomyopathy and characterized by moderate LV dilation and severe systolic dysfunction, together with severe dilation of both atria. Some features of this phenotype are consistent with those reported in other patients carrying the G716R missense mutation [1, 17, 33, 46].

The patient received preventively an automatic defibrillator. Pharmacological management included on-demand diuretics, oral anticoagulants, nonselective β-blockers and angiotensin-converting enzyme (ACE) inhibitors. Due to continued clinical decline to NYHA class IV heart failure, the patient underwent orthotopic heart transplantation at the age of 31. Samples of LV tissue from the explanted heart were utilized in the present study.

### Interstitial fibrosis and cardiomyocytes hypertrophy and disarray in patient left ventricular tissue

Masson-Goldner stained paraffin sections of LV tissue samples showed regions with cellular disarray and fibrosis (collagen-rich fibers) in patient compared with donor tissue (Fig. 2A). Patient’s cardiomyocytes (CMs) were enlarged compared to those from donor (Fig. 2B). This indicates that ventricular hypertrophy occurs mainly due to CMs hypertrophy (also observed in WGA-immunostained transversal sections; Fig. 2D). Yet, large fibrotic regions were found in patient, which favor CMs misalignment (Fig. 2C).

Longitudinal and quasi-transversal sections indicate fewer and smaller fluorescent spots associated to α-actinin and N-cadherin in patient compared to donor tissue (Fig. 2D). This could suggest disturbances of adherent junctions within intercalated discs where CMs join end-to-end along their longitudinal axis of contraction.

Overall, the patients LV myocardium shows quite pronounced hallmarks of HCM even though the expression level of mutant myosin in patient’s LV samples was previously found to be ∼32% G716R-βMyHC [5]. The dissociation between the relatively low mutant protein expression level and myocardial phenotypic severity supports the hypothesis that also for this mutation unequal levels of mutated and wild-type myosin from CM to CM may contribute to development of the HCM phenotype, as shown before for other HCM mutations [23].

### Similar isoform profiles of sarcomeric proteins in mutant and donor myofibrils

Isoform shift in sarcomeric proteins involved in force generation and Ca^2+^-regulation cannot be excluded as a potential adaptive response to HCM. To explore this possibility, sarcomeric proteins were identified in SDS-PAGE gels and western-blots (WBs) loaded with samples from MT-and WT-hvMFs (Fig. 3). G716R mutation is located in the βMyHC isoform (ClinVar:14105) [1]. Our gels analysis indicated that both MT-and WT-hvMFs expressed essentially only the βMyHC isoform, while the αMyHC isoform was not detected (Fig. 3B).

We found both atrial (MLC-1a) and ventricular (MLC-1v) isoforms of the myosin light chain-1 (essential light chain) in MT- and WT-hvMFs. MLC-1v predominated and the small fraction of MLC-1a was similar in both myofibril types (Fig. 3C). Fig. 3D shows that essentially only the ventricular isoform (MLC-2v) of the myosin light chain-2 (regulatory light chain) was expressed in both MT- and WT-hvMFs. No other than the cardiac isoform was detected for the regulatory proteins troponin I (TnI; Fig. 3E) and troponin T (TnT; Fig. 3F). Both α and β tropomyosin (Tm) isoforms were present, with the αTm appearing dominant in both MT- and WT-hvMFs (Fig. 3G).

### G716R mutation influences cross-bridge cycling kinetics without significant effect on force response to different calcium concentrations

Thin bundles of human left-ventricular myofibrils (hvMFs), with sarcomere length (*SL*) of 1.8-2.1 µm, were isolated from the hearts of three healthy donors (WT-hvMFs; Fig. 4A, top-left) and of one patient (MT-hvMFs; Fig. 4A, top-right). Myofibrils were mounted in the setup under relaxing conditions (pCa 8.0).

Depending on the protocol, hvMFs were activated at either saturating (pCa 4.18–4.63) or intermediate (pCa 5.0–6.5) calcium concentrations ([Ca^2+^]) and then relaxed to either very low (pCa 8.0, complete relaxation) or residual (pCa 6.31 or 5.89, partial relaxation) [Ca^2+^]. Prior to Ca^2+^-activation, *SL* was adjusted to 2.3 µm resulting the *F*_pas_, which was subtracted from the total measured force *F*_total_ (Fig. 4A) to calculate the active force *F*_act_ generated by cycling cross-bridges (CBs). Representative force traces illustrating the measured force-related parameters are shown in Fig. 4A,5A.

Regardless pre-incubation with activated catalytic subunit of the protein kinase-A (PKA), at saturating [Ca^2+^] (pCa 4.18), there were no significant differences in *F*_act,max_ between MT-and WT-hvMFs (Fig. 4B; Table S1A). At submaximal Ca^2+^-activation (pCa 5.54; Fig. 4B) without PKA treatment, MT-hvMFs generated higher force than WT-hvMFs as observed also in the normalized *F*_act_–pCa relationships (Fig. 4C). PKA treatment abolished the increased Ca^2+^-sensitivity of force, i.e., pCa_50_ defined as [Ca^2+^] at half-maximal force. Steepness *n*_H_ of the *F*_act_–pCa relationship is interpreted as reflecting the degree of cooperative fast turning-*on*/*off* the troponin-tropomyosin-actin regulatory units along the thin filaments [9]. PKA is often activated during β-adrenergic stimulation of the donor hearts due to inotropic drug administration before organ donation, while the patient received β-blockers. Thus, the potential mutation effect on CB-related force kinetics was further assessed in PKA-treated myofibrils.

Kinetics of Ca^2+^-induced force development are indicated by the activation half-time *t*_1/2,act_ (Fig. 4D; Table S1A,B in *Supplementary Materials*). Rate constants of isometric force redevelopment *k*_tr_ (Fig. 4A,E; 5A-left; Table S1A) were determined after a quick release of isometrically contracting myofibrils to reach essentially zero force and subsequent restretch back to their initial length (Fig. 4A). Upon rapid lowering [Ca^2+^], kinetics of full or partial relaxation are generally described by the relaxation half-time *t*_1/2,rel_ (Fig. 4A,F; Table S1A,B) and more specifically by kinetic parameters *k*_lin_, *t*_lin_, *k*_rel_, detailing the biphasic relaxation (Fig. 5A-right; Table S1A,B). During the first relaxation phase of duration *t*_lin_, sarcomeres remain isometric, while force decays slowly, quasi-linearly with the rate constant *k*_lin_. During the second relaxation phase, sarcomeres lengthen progressively and force decays faster, exponentially with the rate constant *k*_rel_ (>>*k*_lin_), which is influenced by emergent sarcomere-length heterogeneity and compliance (Fig. 4A,5A-right) [19, 41, 42].

Between PKA-treated MT- and WT-hvMFs, there were no major changes in kinetics of Ca^2+^-induced force development at a given pCa (Fig. 4D; Table S1A). The rate constant *k*_tr_, reflecting directly CBs cycling while thin filaments are Ca^2+^-activated, was reduced in MT-vs. WT-hvMFs at pCa 4.18 (Fig. 4E, Fig. 5A-left, Table S1A). The time course for complete relaxation (*t*_1/2,rel_) was significantly prolonged in MT-vs. WT-hvMFs (Fig. 4F, Fig. 5A-right, Table S1A). Different from some other myosin mutations [5, 22], presence of a low level of the mutant G716R-βMyHC seems to influence the sarcomeric ensemble of cycling CBs, affecting particularly myofibrillar relaxation, without increasing force significantly.

To further understand how the heterogeneous ensemble of CBs influence myofibrillar activation and relaxation kinetics, individual rate constants of CB turnover were calculated and used as interpretative tools. In sarcomeres, asynchronous cyclic interactions of myosin heads with thin filaments during force production can be classified into two groups of functional states: force-generating (FG) and non-force-generating (nFG) states (Fig. 5A-left) [7]. In FG-states, heads are strongly bound to actin and proceed with the power-stroke, while in nFG-states they bind weakly to actin with rapid attachment/detachment kinetics. When isometric force is generated, participating CBs are cycling between FG- and nFG-states. The apparent rate constant *f*_app_ represents the probability of CB transition from nFG- to FG-states (mainly associated with the steps related to P_i_ release during the ATP-turnover), while *g*_app_ represents the transition back to nFG-states (associated with ADP release and ATP-induced myosin-actin dissociation) (Fig. 5A-left). *k*_tr_ (=*f*_app_+*g*_app_) reflects kinetics of CBs re-distribution between FG- and nFG-states. *f*_app_ is influenced by [Ca^2+^], because it depends on the Ca^2+^-regulated fast equilibrium between turned-*on/*-*off* regulatory units along thin filaments. *g*_app_, which has been shown to correspond to *k*_lin_ and to modulate tension cost (=ATPase activity/isometric force), is independent on [Ca^2+^] [7, 9, 19]. At different [Ca^2+^], isometric force is primarily modulated by the duty-ratio (i.e., the fractional occupancy of cycling myosin interacting strongly vs. weakly with thin filaments) being quantified by the *turnover-kinetic-ratio f*_app_/(*f*_app_+*g*_app_). Force is therefore defined as *F*_act_=*N*_r_·*F*_0_·*f*_app_/(*f*_app_+*g*_app_) [7]. *N*_r_ is the number of recruited, actively turning-over myosin heads per half-sarcomere. *F*_0_ is the mean force generated by a single CB in the FG-states.

Calculations based on measured values for *k*_tr_ and *k*_lin_ show that *f*_app_ (=*k*_tr_-*k*_lin_) and duty-ratio *f*_app_/(*f*_app_+*g*_app_) were significantly reduced in MT-hvMFs compared with WT-hvMFs at both pCa 4.18 and pCa 5.54, while *g*_app_ remained unaffected by the mutation (Fig. 5B). Therefore, similar isometric forces *F*_act_ generated by patient and donor myofibrils (Fig. 4B,C) suggest that the *ensemble-force-factor N*_r_·*F*_0_ can be higher in MT-hvMFs, containing both MT- and WT-CBs, than in WT-hvMFs.

### Residual calcium concentration exacerbates relaxation dysfunction

Previous studies investigating mechanisms of HCM at the cellular level revealed that in patient’s CMs, the intracellular [Ca^2+^]_i_ can be higher during diastole than in healthy CMs [11]. hvMFs analyzed in the present study were isolated form frozen patient or donor tissues, from which preparation of viable CMs was not feasible. Nevertheless, since the patient developed diastolic dysfunction, which could partially be caused also by increased intracellular diastolic [Ca^2+^]_i_ in the HCM-patient’s hypertrophic CMs, we aimed to test this by analysis of relaxation kinetics at residual Ca^2+^-levels. The protocol was adjusted so that fully Ca^2+^-activated (pCa 4.63) myofibrils were partially relaxed either at pCa 6.31 or pCa 5.89. Results were compared with those of completely relaxed hvMFs at pCa 8.0, as well as between MT-and WT-hvMFs at each pCa to which myofibrils were relaxed (Fig. 6).

At pCa 6.31 and pCa 5.89, partially relaxed (PKA-treated) MT- and WT-hvMFs generated similar submaximal isometric forces (Fig. 6B). There were no significant differences in *k*_lin_ and *t*_lin_ parameters between MT- and WT-hvMFs at each pCa, except for *t*_lin_ at pCa 8.0 (Fig. 6C,D; Table S1 in *Supplementary Materials*). *k*_rel_ was significantly reduced in relaxing MT-vs. WT-hvMFs at all pCa-values (Fig. 6E; Table S1A,B). *t*_1/2,rel_ was significantly longer for MT-compared to WT-hvMFs during relaxation at all pCa-values (Fig. 6F; Table S1A,B). Assuming that diastolic intracellular [Ca²⁺]ᵢ is elevated only in patient’s CMs, impairment of relaxation kinetics becomes even more pronounced when partially relaxed MT-hvMFs are compared with fully relaxed donor WT-hvMFs (Fig.6D,E,F). This could well represent a sarcomeric contribution to diastolic dysfunction in the patient.

### Impairment of mutant myofibrils relaxation is worsened by mavacamten

Mavacamten (mava) was designed to normalize hypercontractile function of the myocardium and was recently approved for treatment of HOCM patients [6, 32]. We have previously shown that the myosin ATPase inhibitor BDM accelerated force decay, diminishing the difference in relaxation kinetics between control and mutant ventricular myofibrils, in which the HCM-related ΔK184-cTnI mutation impaired the relaxation [18]. It was also shown that 0.5 µM mavacamten accelerated relaxation kinetics of donor WT-hvMFs [36]. In this line of argumentation, we hypothesized that the impairment in relaxation kinetics of MT-hvMFs will be diminished by mavacamten. Therefore, we compared force-related parameters of PKA-treated MT-and WT-hvMFs in absence (-mava) or presence (+mava) of 0.6 µM mavacamten (IC_50_=0.58±0.07≍0.6 µM for hvMFs; [36]) (Fig. 7). The results of MT-hvMFs in presence of the drug (+mava) can also be compared with those of WT-hvMFs in absence of the drug, assuming that only the HOCM-patient receives the mavacamten.

Isometric force (*F*_act_) generated by WT-hvMFs was reduced by mavacamten at submaximal [Ca^2+^] (-89% at pCa 5.89) more than at saturating [Ca^2+^] (-40% at pCa 4.63) (Fig. 7A). This suggests that drug’s inhibitory effect depends on [Ca²⁺] at which myofibrillar force is generated. In MT-hvMFs, inhibition of *F*_act_ by mavacamten was opposite than in WT-hvMFs and less efficient at partial [Ca^2+^]: -27% at pCa 5.89 and -50% at pCa 4.63 (Fig. 7A). Therefore, in patient’s myofibrils, the drug’s potency might be weakened at physiological submaximal [Ca^2+^].

Kinetics of Ca^2+^-induced maximum force development were significantly slowed down by mavacamten in both myofibril categories (larger activation half-time *t*_1/2,act_; Fig. 7B), with a less pronounced effect on MT-hvMFs. *k*_tr_ at saturating [Ca^2+^] of WT-hvMFs was slowed down substantially (-32%) with the drug, but its effect was much smaller (-13%) in MT-hvMFs (Fig. 7C). Thus, the mutation-induced reduction (-23%) in *k*_tr_ of MT-hvMFs was further reduced (-32%) by mavacamten at pCa 4.63.

WT-hvMFs full relaxation at pCa 8.0 was accelerated by mavacamten as indicated by significantly reduced (-34%) relaxation half-time *t*_1/2,rel_ (Fig. 7D). This reinforces previous findings that the myosin inhibitor accelerates myofibrillar relaxation kinetics. In the absence of mavacamten (Fig. 4F,6F), *t*_1/2,rel_ of MT-hvMFs at full (pCa 8.0) and partial (pCa 5.89) relaxation was significantly prolonged compared to fully relaxed WT-hvMFs (Fig. 7D). This likely corresponds to the mutation effect. Yet, when MT-hvMFs were treated with mavacamten, *t*_1/2,rel_ was not reduced as expected, and as observed in WT-hvMFs, and the mutation-induced impairment was even more pronounced at partial relaxation (Fig. 7D). Thus, mimicking the possibility of partial relaxation in HCM myocardium with MT-hvMFs, mavacamten seems to substantially exacerbate the impairment of myofibrillar relaxation kinetics.

Looking at the myofibrillar relaxation in more detail, the rate constant *k*_lin_ of the first phase, in the absence of mavacamten, was similar for both MT-and WT-hvMFs at full (pCa 8.0) and partial (pCa 5.89) relaxation, respectively (Fig. 6C,7E). At full relaxation, the drug increased *k*_lin_ in WT-hvMFs (+42%) and in MT-hvMFs (+47%) when compared with WT-hvMFs in absence of mavacamten (Fig. 7E). In contrast, kinetics of the first phase at pCa 5.89 in MT-hvMFs was drastically reduced (-32%) with mavacamten (Fig. 7E). The duration of the first relaxation phase, *t*_lin_ was slightly increased (+16%) in MT-vs. WT-hvMFs only at full relaxation (Fig. 6D,7F). Mavacamten had no significant effect on *t*_lin_ of WT-hvMFs. Yet, in MT-hvMFs, the drug increased *t*_lin_ by +31% during full relaxation and by +114% at partial relaxation (Fig. 7F), which is consistent with slower *k*_lin_. Thus, the drug delayed the onset of the second relaxation phase in MT-hvMFs.

The rate constant *k*_rel_ of the second relaxation phase of MT-hvMFs was reduced at both full (-29%) and partial (-62%) relaxations compared to fully relaxed WT-hvMFs (Fig. 7G, cf. also Fig. 6E), likely indicating the mutation effect. In WT-hvMFs only at full relaxation, mavacamten substantially increased *k*_rel_ (+102%; Fig. 7G). Notably, in MT-hvMFs at full relaxation (-39%), and even more at partial relaxation (-73%), the drug further reduced *k*_rel_ when compared with WT-hvMFs at full relaxation in absence of mavacamten (Fig. 7G), thus aggravating the impairment of relaxation in patient’s MT-hvMFs.

In summary, when compared with baseline WT-hvMFs, mavacamten exacerbated the detrimental functional effects of the mutation in MT-hvMFs. It reduced isometric force at saturating and intermediate [Ca^2+^], and slowed *k*_tr_ even more. Most importantly, mavacamten did not compensate the relaxation impairment and further slowed down relaxation parameters at residual calcium levels in MT-hvMFs.

Calculations of the individual rate constants for CB turnover, *f*_app,max_ and *g*_app_, showed that mavacamten reduces *f*_app,max_ (-69%) and increases *g*_app_ (+42%) in WT-hvMFs, but less in MT-hvMFs (Fig. 8A,B). Compared to WT-hvMFs in absence of mavacamten, the mutation-related reduction (-48%) of *f*_app,max_ was further lowered (-60%) by the drug (Fig. 8A). Consequently, mavacamten reduced (-51%) the duty-ratio *f*_app,max_/(*f*_app,max_+*g*_app_) in WT-hvMFs, but interestingly, much less in MT-hvMFs (Fig. 8C).

The *ensemble-force-factor N*_r_·*F*_0_ (Fig. 8D) can be calculated from *F*_act,max_ (Fig. 7A) and duty-ratio (Fig. 8C). In the absence of mavacamten, *N*_r_·*F*_0_ for MT-hvMFs (with the mixed population of MT-CBs and WT-CBs) was significantly higher (+34%) compared to WT-hvMFs (Fig. 8D). Upon mavacamten (0.6 µM) exposure, *N*_r_·*F*_0_ decreased slightly (-14%) in WT-hvMFs, but substantially (-43%) in MT-hvMFs, and less (-23%) when compared with WT-hvMFs in absence of the drug.

## DISCUSSION

We aimed to understand functional consequences of the HCM-associated heterozygous β-myosin G716R missense mutation in absence and presence of mavacamten in patient LV myofibrils (MT-hvMFs) compared with donor myofibrils (WT-hvMFs) as control. Although previous studies indicated that several HCM-mutations caused hyperfunction in investigated contractile models [40], our results showed that sarcomeric isometric force and its sensitivity to varying [Ca^2+^] were not significantly altered in MT-hvMFs containing a mixture of ∼32% MT-and 68% WT-myosin heads [5]. Nevertheless, G716R-βMyHC impaired maximum rate constants of myofibrillar force generation and relaxation kinetics compared to WT-hvMFs. The most notable finding is that, while the myosin inhibitor mavacamten operated as expected in donor myofibrils and accelerated relaxation kinetics, its functional effect on patient myofibrils during relaxation was the opposite.

### Illustrative βMyHC structural models indicate locations of the mutation and mavacamten binding

G716 (Fig. 1A,B) lies adjacent to a hydrophobic pocket defined by L700, I736, and L489, which forms a conformational-sensitive pivot within the converter domain and relay helix [3, 4, 29]. This region may become surface-exposed as the converter rotates in the active/pre-power-stroke state [4]. The small, neutral residue G716 is conserved across mammalian myosin. Introducing the positively charged bulky chain residue R716 (Fig. 1E) could weaken the local hydrophobic stabilization and may impose steric restrictions on the converter’s surface topology.

Mavacamten binds to an allosteric hydrophobic pocket (Fig. 1) that bridges the catalytic domain, converter, and relay regions interacting with I713, R721 and L770 [3, 29]. Depending on the R716 rotamer, the substitution G716R may alter the local electrostatic landscape and create direct steric crowding with either the drug or the flanking I713 and R721 long residues, thus reducing local flexibility. The potential point mutation-induced modifications in the energetic landscape of the motor may alter converter geometry exposing different local binding options to mavacamten [3]. This may further influence the transition towards the power-stroke state which could be reflected in cross-bridge cycling contributing to force generation, as we observed here in myofibrils. It could also alter the equilibrium between *on*- and *off*-states of myosin heads in a different way than in absence of the drug, as we can indirectly infer from our analysis and discuss within the frame of previously published studies. Further biochemical, structural and computational studies could test these hypotheses.

### Mutation-induced functional impact under isometric conditions

Force is modulated kinetically by the duty-ratio *f*_app_*/(f*_app_+*g*_app_) [7], as calcium-dependent thin-filament regulation rapidly controls the fraction of myosin heads in force-generating states (Fig. 5A, inset), while they are cycling asynchronously between weak (nFG-states) and strong (FG-states) actin binding conformations. In a wild-type contractile model, these events occur during ATP-turnover on a shorter timescale than that considered for the equilibrium between myosin *off*↔*on* states interconversion [30, 44].

We hypothesized that the G716R substitution in βMyHC perturbs cycling CBs in myofibrils from the heterozygous patient with sarcomeres containing a mixture of MT- and WT-βMyHC [5].

In MT-vs. WT-hvMFs, the duty-ratio of cycling CBs is lower because of reduced *f*_app,max_, which seems to be more sensitive to the mutation than *g*_app_ (Fig. 5B,7H-J). Impaired *f*_app,max_ can be associated with slower P_i_-release, which may favor accumulation of weak acto-myosin A∼M_MT_·ADP·P_i_ rather than A∼M_WT_·ADP·P_i_ intermediates during steady-state ATP-turnover of the heterogeneous CB ensemble in MT-hvMFs. Thus, when dimeric heads are in the pre-power-stroke conformation, which was considered to favor IHM formation [30], IHM with MT/WT-and MT/MT-heads may have a shorter lifetime than IHM with WT/WT-heads.

As *F*_act_=*N*_r_·*F*_0_·*f*_app_/(*f*_app_+*g*_app_), similar isometric force *F*_act_ generated by MT-and WT-hvMFs (Fig. 4B,7A) and the reduced duty-ratio *f*_app_/(*f*_app_+*g*_app_) in MT-compared with WT-hvMFs (Fig. 5B,8C) implies that the *ensemble-force-factor N*_r_·*F*_0_ is significantly augmented in MT-hvMFs compared to WT-hvMFs (Fig. 8D). This suggests that binding of the drug to myosin in a region near the G716R substitution may reduce the number of turning over myosin heads (decreasing *N*_r_) or reduce specific *F*_0_, while both effects could also occur.

Assuming that allosteric conformational change induced by the HCM-mutation favors myosin’s *on*-state by disrupting the IHM configuration [34, 37, 40], more MT-heads and WT-heads from the pool of MT/WT-and MT/MT-myosin dimers in MT-hvMFs may become available to cycle within the nFG↔FG equilibrium. Thus, an increase in the number (*N*_r_=*N*_r,MT_+*N*_r,WT_) of recruited heads would be expected and this could cause a net increase of the estimated *ensemble-force-factor N*_r_·*F*_0_, as calculated here (Fig. 8D). If *F*_0_ of MT-heads would be reduced compared to *F*_0_ of WT-heads, as suggested with *Dictyostelium* myosin expressing the mutation at an equivalent position [12], newly available WT-heads (in *on*-state) from the MT/WT-myosin dimers could compensate and even cause a net increase of the *ensemble-force-factor N*_r_·*F*_0_. Other HCM-associated R723G and R719W mutations in converter domain were previously shown rather to increase *F*_0_ per CB in the FG-states [22, 38]. While Arg in G716R substitution acts primarily via steric crowding and charge insertion, Gly in R723G introduces excessive local backbone flexibility decreasing the electrical charge. The rigid aromatic indole ring of Trp in R719W drastically alters the surface potential from charged hydrophilic/basic to hydrophobic. Consequently, these HCM-related mutations (G716R, R723G, R719W) could distinctly influence allosteric changes in myosin. Therefore, the opposite effect in *F*_0_ reported with *Dictyostelium* myosin analog [12] may not be automatically unexpected, even if all three HCM-mutations are in converter domain.

In the sarcomeric MT/WT-CBs ensemble, the mutation may affect both the cycling CB nFG↔FG and myosin-head *off*↔*on* equilibria operating on different time-scales. Their interplay, which may depend on the fraction of MT-CBs in sarcomere, could in turn modulate kinetics of myofibrillar force response and thus propagate through mechano-sensing pathways to upstream cellular responses. This mechanism may contribute, along with other determinants, to cause a net sarcomeric hypo-, normo-or hypercontractile phenotype.

### Functional effects of mavacamten under isometric conditions

Previous studies concluded that mavacamten reduced force by decreasing *N*_r_ of CBs participating in force generation [2, 31, 35]. It also slows P_i_-release kinetics, inhibiting steady-state ATP-turnover, and favoring accumulation of low-force, pre-power stroke A∼M·ADP·P_i_ cross-bridges conformation [21, 31].

In this study, the drug at partial Ca^2+^-activation inhibited force less in MT-than in WT-hvMFs, while at maximum activation inhibition was comparable (Fig. 7A). This suggests that the inhibitory effect of mavacamten in MT-hvMFs may depend on the equilibrium of thin filament-mediated Ca^2+^-regulation of cycling CBs reflected by *f*_app_ and *f*_app_/(*f*_app_+*g*_app_) duty-ratio. In maximally Ca^2+^-activated MT-hvMFs, these parameters were less reduced by the drug than in WT-hvMFs (Fig. 8A,C). The initial decreased *f*_app,max_ in MT-vs. WT-hvMFs was further diminished by mavacamten in MT-hvMFs (Fig. 8A). The rather small effect of mavacamten on *f*_app,max_ and duty-ratio of MT/WT-CBs ensemble in MT-hvMFs suggests that binding of the drug to MT-heads might occur differently than to WT-heads, likely due to the proximity of the G716R-mutation to the mavacamten binding region in myosin [3]. This may alter allosteric molecular changes which could further influence pre-power stroke and power-stroke states during the CB cycle.

G716R induced an increase of the *ensemble-force-factor N*_r_·*F*_0_ (Fig. 8D). In WT-hvMFs, mavacamten seems to have a reduced effect on *N*_r_·*F*_0_ arising from sequestration of WT-heads in the *off*-state [2]. Yet, a substantial inhibition likely occurs through reducing *f*_app,max_ and duty-ratio with mavacamten. Therefore, in MT-hvMFs with a mixed population of MT-and WT-heads, the larger mavacamten-induced decrease in *N*_r_·*F*_0_ (where *N*_r_=*N*_r,MT_+*N*_r,WT_) might essentially be due to a substantial reduction of the *ensemble-force-factor* of the MT-CBs. Mavacamten may alter the nano-environment induced by G716R, favoring MT-CBs to populate weak acto-myosin states mainly by reducing *f*_app,max_ rather than increasing *g*_app_ (Fig. 8A,B) and increasing population of MT-heads in the *off*-state, assuming that *F*_0_ in MT-vs. WT-CBs is either reduced or not altered (Fig. 8D).

### Relaxation impairment in mutant myofibrils

Impaired relaxation (Fig. 4F,6F,7D) may reflect an intrinsic effect of the myosin motor mutation, as the similar isometric force responses of MT-and WT-hvMFs at low Ca²⁺ concentrations (Fig. 4B,C,6B) indicate no apparent evidence of residual bound CBs at pCa 8.0, a phenomenon reported for some other HCM mutations [18].

Previous studies revealed that diastolic intracellular [Ca^2+^]_i_ in HCM cardiomyocytes (CMs) can be higher than in WT-CMs [11]. Assuming this scenario in CMs of the HCM-patient exhibiting diastolic dysfunction, Ca^2+^-activated myofibrils at pCa 4.63 were relaxed either fully at pCa 8.0 or partially at pCa 5.89 or pCa 6.31. These experiments showed that at residual [Ca^2+^], biphasic relaxation impairment of MT-hvMFs was exacerbated (Figs. 6A,F,7D).

### First relaxation phase

Single molecule studies indicated that CB detachment is load dependent and elevated resisting load prolongs the lifetime of strong CBs [25]. During the first phase of isometric relaxation, attached CBs can be thought of as detaching under relatively high mechanical pre-load contributing to the rate constant *g*_app_ (=*k*_lin_) of the CBs ensemble leaving FG-states [41, 42], which appears unaffected by MT-CBs in MT-vs. WT-hvMFs (Fig. 5B,6C,7E). At full relaxation, the small increase in *t*_lin_ of MT-hvMFs was further augmented at partial relaxations when compared with WT-hvMFs relaxing at pCa 8.0 (Fig. 6D,7F). This augmentation of *t*_lin_ at partial relaxation might be due to few additional myosin heads populating the FG-states [43].

### Second relaxation phase

Once a sarcomere starts to lengthen at the onset of the second phase, its elongation reduces the average strain in neighboring sarcomeres. *k*_rel_ (>>*k*_lin_) is dominated by accelerated detachment of the remaining CBs as local pre-load progressively diminishes [41, 42]. Reduced pre-load per CB implies lower mechanical stress on the converter domain (where G716R is located), which is a major determinant of CB elasticity [22, 38]. In MT-compared with WT-hvMFs, *k*_rel_ was significantly slower under all relaxation conditions (Fig. 6E,7G), indicating impaired relaxation kinetics in patient myofibrils containing MT-CBs that leave FG-states while moderately pre-loaded. This finding is consistent with the reduced sliding velocity of unloaded actin filaments driven by analogue MT-vs. WT-myosin heads [12].

This interpretation may further hypothesize about how mechanical load could influence cycling CBs during steady-state on the timescale of ATP-turnover. When sarcomeres contract against a moderate load, the effects of G716R substitution suggest nano-structural alterations in the converter domain which may result in prolonged lifetime of the A-M_MT_·ADP conformation slowing ADP-release. On contrary, when sarcomeres contract against an elevated load, the mutation may only minimally interfere with the A-M_MT_·ADP conformation. This hypothesis could further indicate that in the ensemble of cycling MT/WT-CBs in MT-hvMFs, the threshold of MT-CBs for a functional consequence may depend on the mechanical preload experienced by cycling CBs in sarcomeric environment.

### Functional effects of mavacamten in mutant myofibrils during relaxation

Unexpectedly, mavacamten did not rescue the impaired relaxation of MT-hvMFs.

### First relaxation phase

As previously reported [36], *k*_lin_ was accelerated significantly by mavacamten in WT-hvMFs (Fig. 7E), probably due to reducing the lifetime of the A-M_WT_·ADP conformation of strongly-bound actin-myosin prior to leaving FG-states. This effect seems diminished in fully relaxing MT-hvMFs and even reversed at partial relaxation (Fig. 7E), while *t*_lin_ was substantially prolonged (Fig. 7F). This suggests that mutation may prevent mavacamten-induced acceleration effect of *g*_app_ (which reflects the probability of CBs leaving FG-states) in the sarcomeric highly pre-loaded MT/WT-CBs ensemble.

### Second relaxation phase

At full relaxation of WT-hvMFs, mavacamten increased *k*_rel_, as previously observed [36], by accelerating remaining moderately pre-loaded WT-CBs leaving FG-states (Fig. 7G). During partial relaxation of WT-hvMFs, the presence of residual force-generating CBs seems to blunt acceleration induced by mavacamten (Fig. 7G).

When compared with fully relaxing WT-hvMFs in absence of mavacamten, *k*_rel_ of MT-hvMFs was further reduced with the drug during both, full and partial relaxation, suggesting prolonged lifetime of the A-M_MT_·ADP conformation with the drug bound to moderately pre-loaded MT-heads (Fig. 7G).

The G716R-mutation effects, together with the effects of mavacamten and residual CBs during partial relaxation, could also explain the large delay (*t*_lin_) of the onset of the second relaxation phase in MT-hvMFs. Prolonged *t*_lin_ and the strong reduction of *k*_rel_ at partial relaxation contributes to the notable large increase of the overall relaxation half-time *t*_1/2,rel_ of MT-hvMFs in the presence of the drug (Fig. 7D).

### Sarcomeric dysfunction convolutes with histopathological alterations

Replacement interstitial fibrosis (Fig. 2) increases myocardial stiffness leading to elevated LV end-diastolic filling pressure. This, together with impairment of myofibrils relaxation, including the possibility of moderate increase in intracellular diastolic [Ca^2+^]_i_, could aggravate patient’s diastolic dysfunction.

Cardiomyocyte disarray, interstitial fibrosis [15] and spatially impaired N-cadherin-mediated coupling [10] may decrease intercellular and extracellular-matrix-mediated force transmission, potentially compromising the mechanical coherence of the myocardium. Force vectors of individual CMs can be oriented in different directions [24]. Cell-to-cell variability in mutation expression, likely causing inherent variability in contractile performance cannot be excluded [23]. Consequently, the net force output in the direction of contraction may decrease, leading to compromised regional or global ventricular function.

Further investigations could validate a potential disturbance in expression of N-cadherin which may impact mechano-transduction and contribute to fibrosis. This may represent a signature of the disease transition from hypertrophic to dilative phenotype and heart failure, explaining reduction of the LV obstruction with progression of the disease. Overall, multiscale heterogeneity may underlie the histological abnormalities and aligns with clinical observations of impaired systolic and diastolic function in this patient.

### Limitations of the study

Only MT- and WT-hvMFs thin bundles meeting predefined quality criteria were investigated: diameter 1-4µm, length 10-50 µm, generally cylindrical shape, preserved sarcomeric pattern, and minimal sarcomere length heterogeneity. Compared to the investigated donor myofibrils, small deviations from these criteria were sometimes observed in 10-20% patient myofibrils, which were not investigated. This suggests that some functional differences of patient vs. control myofibrils could be slightly underestimated.

In general, measurements with myofibrils typically exhibit greater variability than studies using other myocardial preparations. To minimize potential myofibrillar force run-down, thin myofibrillar bundles were subjected to three successive Ca^2+^-activations under routine experimental conditions. For force-pCa measurements, the protocol was extended to seven activations to obtain the complete Ca²⁺-dependence of force development preserving myofibrillar contractile stability.

While control myofibrils were available from three donor hearts, myofibrils containing the G716R mutation in myosin were isolated from only one available explanted heart.

## Conclusion

The present investigation should be regarded as a mechanistic study mainly addressing specific aspects of cross-bridge function within the sarcomere, rather than as an evaluation of clinical outcomes or therapeutic efficacy.

This study suggests that HCM-related β-myosin mutation G716R does not evoke gain-of-function in human left ventricular sarcomeres containing a heterogeneous population of mutant and wild-type cross-bridges. G716R-βMyHC reduced the maximum probability of the cross-bridges to enter force-generating states and substantially impaired myofibrillar relaxation kinetics. Unexpectedly, mavacamten failed to restore the impaired relaxation of G716R-patient-derived myofibrils, in contrast to the effect of the drug on relaxation of WT-myofibrils from donor heart. This indicates that mutant cross-bridges exhibit a differential response to mavacamten compared with wild-type cross-bridges. The results suggest that if patient’s cardiomyocytes would exhibit persistently elevated residual diastolic Ca²⁺ concentrations, mavacamten may further aggravate the diastolic deficit.

Although mavacamten targets cardiac sarcomeric β-myosin, its effects extend to whole-heart function, with demonstrated clinical benefit in many HOCM-patients [28]. Nevertheless, the mechanisms underlying inter-individual variability in treatment response remain incompletely understood. Our findings suggest, but do not establish, that HCM-associated mutations may modulate myofibrillar sensitivity to mavacamten, potentially informing more individualized treatment strategies.

## ACKNOWLEDGMENTS

We wish to acknowledge the patient, donors and all persons involved in administrating the tissue procurement.

The authors thank to Dr. Meißner Joachim for the valuable feedback on the manuscript and to Alexander Lingk, Torsten Beier and Uwe Krumm (Molecular and Cell Physiology, Hannover Medical School) for their excellent technical assistance.

We gratefully acknowledge the support of the Research Core Unit for Laser Microscopy at Hannover Medical School, as well as the acquisition of histological images using the Metafer scanning system facilitated by Prof. Dr. J.H. Bräsen (Nephropathology Unit, Institute of Pathology, Hannover Medical School).

## AUTHOR CONTRIBUTIONS

Conceptualization and project design: BI

Methodology: BI, BP, TH, CR

Investigations: AB, BP, TH, AF

Data analysis: AB, BP, TH, BI

Supervision: BI, TK

Writing – original draft: BI

Writing – review & editing: BI, AF, TK

All authors approved the final version of the manuscript.

## SUPPLEMENTARY MATERIALS

### RESULTS

**Table S1.** Functional parameters of patient and donor ventricular myofibrils. Steady-state and kinetic force-related parameters resulting from isometrically contracting-relaxing human left ventricular myofibrils (hvMFs) isolated from patient (MT-hvMFs) and donor (WT-hvMFs) tissues. Myofibrils were activated either at saturating (pCa 4.18) or at partial (pCa 5.54) calcium concentrations, as indicated for parameters related to force generation (*F*_act_, *t*_1/2,act_, *k*_tr_). Relaxation kinetic parameters (*k*_lin_, *t*_lin_, *k*_rel_, *t*_1/2,rel_) were derived from initially fully Ca^2+^-activated myofibrils which were then relaxed either at pCa 8.0 (full relaxation) or at pCa 6.31 or pCa 5.89 (partial relaxation). Values are given as mean±SEM; n is the number of myofibrils contributing to the indicated force-related parameter; WT-hvMFs were prepared from a total of 3 different donor hearts (data is pooled); significance indicated by # (Student’s *t*-test) of the differences between the mean values of the corresponding parameters (MT-hvMFs vs. WT-hvMFs) is indicated by the *p* value.

| **Force-related parameters** | **Before treatment with protein kinase A (-PKA)** |  |  |  |  | **After treatment with protein kinase A (+PKA)** |  |  |  |  |
|---|---|---|---|---|---|---|---|---|---|---|
|  | **WT-hvMFs** | **n<sub>WT</sub>** | **MT-hvMFs** | **n<sub>MT</sub>** | ***p*** | **WT-hvMFs** | **n<sub>WT</sub>** | **MT-hvMFs** | **n<sub>MT</sub>** | ***p*** |
| **A.** |  |  |  |  |  | ***Calcium-induced activation*** |  |  |  |  |
| ***F*<sub>act,max at pCa 4.18</sub> (kPa)** | 53±2 | 93 | 53±6 | 33 | 0.7178 | 47±4 | 35 | 55±2 | 36 | 0.0503 |
| ***F*<sub>act at pCa 5.54</sub> (kPa)** | 26±1 | 68 | 36±1<sup>#</sup> | 33 | <0.0001 | 24±1 | 30 | 28±2 | 18 | 0.0562 |
| **pCa<sub>50</sub>** | 5.56±0.02 | 68 | 5.74±0.03<sup>#</sup> | 33 | <0.0001 | 5.53±0.02 | 30 | 5.54±0.02 | 18 | 0.7295 |
| ***n*<sub>H</sub>** | 1.69±0.06 | 68 | 1.59±0.10 | 33 | 0.3797 | 2.06±0.11 | 30 | 2.22±0.13 | 18 | 0.3604 |
| ***t*<sub>1/2,act at pCa 4.18</sub> (ms)** | 1179±50 | 68 | 1330±66 | 30 | 0.0844 | 1307±69 | 29 | 1517±85 | 16 | 0.0684 |
| ***t*<sub>1/2,act at pCa 5.54</sub> (ms)** | 2947±118 | 50 | 2950±153 | 31 | 0.9902 | 3479±270 | 23 | 3953±400 | 18 | 0.3164 |
| ***k*<sub>tr,max</sub> (s<sup>-1</sup>)** | 0.62±0.02 | 85 | 0.55±0.03 | 29 | 0.0781 | 0.59±0.03 | 30 | 0.48±0.02<sup>#</sup> | 38 | 0.0013 |
| ***k*<sub>tr at pCa 5.54</sub> (s<sup>-1</sup>)** | 0.35±0.01 | 56 | 0.37±0.02 | 22 | 0.3242 | 0.32±0.01 | 26 | 0.27±0.02 | 17 | 0.0690 |
|  |  |  |  |  |  | ***Full relaxation at pCa 8.0*** |  |  |  |  |
| ***k*<sub>lin</sub> (s<sup>-1</sup>)** | 0.22±0.01 | 58 | 0.23±0.02 | 24 | 0.6851 | 0.21±0.01 | 65 | 0.22±0.02 | 35 | 0.6314 |
| ***t*<sub>lin</sub> (ms)** | 221±10 | 60 | 240±11 | 33 | 0.2052 | 214±10 | 33 | 267±18<sup>#</sup> | 26 | 0.0087 |
| ***k*<sub>rel</sub> (s<sup>-1</sup>)** | 3.28±0.13 | 62 | 2.90±0.15 | 33 | 0.0783 | 3.42±0.12 | 33 | 2.94±0.15<sup>#</sup> | 26 | 0.0126 |
| ***t*<sub>1/2,rel</sub> (ms)** | 436±13 | 54 | 493±14<sup>#</sup> | 22 | 0.0140 | 418±9 | 33 | 530±16<sup>#</sup> | 26 | <0.0001 |
| **B.** |  |  |  |  |  | ***Partial relaxation at pCa 6.31*** |  |  |  |  |
| ***k*<sub>lin</sub> (s<sup>-1</sup>)** |  |  |  |  |  | 0.17±0.02 | 30 | 0.21±0.02 | 19 | 0.1304 |
| ***t*<sub>lin</sub> (ms)** |  |  |  |  |  | 388±27 | 38 | 362±29 | 19 | 0.5629 |
| ***k*<sub>rel</sub> (s<sup>-1</sup>)** |  |  |  |  |  | 1.99±0.11 | 38 | 1.17±0.07<sup>#</sup> | 19 | <0.0001 |
| ***t*<sub>1/2,rel</sub> (ms)** |  |  |  |  |  | 751±52 | 30 | 915±47<sup>#</sup> | 19 | 0.0357 |
|  |  |  |  |  |  | ***Partial relaxation at pCa 5.89*** |  |  |  |  |
| ***k*<sub>lin</sub> (s<sup>-1</sup>)** |  |  |  |  |  | 0.18±0.02 | 42 | 0.23±0.01 | 19 | 0.0525 |
| ***t*<sub>lin</sub> (ms)** |  |  |  |  |  | 376±29 | 53 | 432±35 | 19 | 0.2936 |
| ***k*<sub>rel</sub> (s<sup>-1</sup>)** |  |  |  |  |  | 2.60±0.18 | 53 | 1.31±0.08<sup>#</sup> | 19 | <0.0001 |
| ***t*<sub>1/2,rel</sub> (ms)** |  |  |  |  |  | 695±49 | 40 | 912±49<sup>#</sup> | 19 | 0.0075 |

### MATERIALS and METHODS

#### Patient data

*Ethical statement*: Informed consent was obtained from all individuals according to approved ethics committee protocols of the institutions involved. All experiments performed with samples of human cardiac tissue (Vote No. 2276/2014) and human skeletal muscle tissue (Vote No. 2729/2001) were approved by the Ethics Committee of Hannover Medical School (MHH). Approval of MHH Ethics Committee was obtained for anonymized use of the different muscle samples in our study. The investigations conformed to the principles outlined in the Declaration of Helsinki (World Medical Association, 1997). All subjects gave written informed consent at the different institutions.

Human cardiac tissue samples from the patient were provided from Hospital Clinic, Cardiology Department, Barcelona. Myocardial tissue was obtained upon cardiac transplant from the left ventricle (LV) of a 32 years old male patient (Id: M030/H92) and preserved frozen at -196°C in liquid nitrogen. Sample sequencing revealed the heterozygous HCM-related missense point mutation in the *MYH7* gene (c.2146G>A) encoding for βMyHC (G716R). Additional information regarding a patient carrying the Gly716R mutation can be found in Annan et al., 1994 [1].

Control tissue was from 3 non-transplanted donor hearts (all males) for which no suitable recipient was found: H105 sample (Id: 5.105; donor age 65), H107 sample (Id: 5.133; donor age 48), H108 sample (Id: 6.004; donor age 48). These samples were provided by St. Vincent’s Hospital in Sydney, Australia, the Sydney Heart Bank [3] and preserved frozen at -196°C in liquid nitrogen. The donor hearts had no known impairment of cardiac function. There was no mutation within this gene except for a few single nucleotide polymorphisms (SNPs): H105 had three SNPs (c.732C>T; c.2967T>C; c.5106G>A) and H108 had one SNP (c.5106G>A) within the *MYH7* gene [8], and they were considered as wild-type (WT) regarding known HCM/DCM/RCM-related gene mutations in *MYH7*. Our previous studies showed no age-dependent differences in any of the parameters of sarcomere function in control samples [5].

As reference for protein analysis, we also used tissues (provided from MHH) from human atrium (female, age 76; Id: hAD6) and human gastrocnemius (male, age 19; Id: H14/15).

#### Solutions composition

*Na-solution* (ionic strength 206.5 mM, pH 7.0 at 5°C, pCa 8.0) contains: 5 mM KH_2_PO_4_ (Fluka), 5 mM Mg-acetate (Sigma), 5 mM EGTA (Sigma), 3 mM Na_2_ATP (Merck), 50 mM NaCrP (Sigma) and 8 mM NaN_3_ (Serva), 5 mM dithiothreitol (DTT; Sigma) and protease inhibitory cocktail (PIC; Sigma). *K-solution* (ionic strength 206.5 mM, pH 7.0 at 5°C, pCa 8.0) contains: 10 mM Imidazole (Sigma), 139.6 mM KCl (Merck), 6.04 mM MgCl_2_ (Sigma), 2 mM EGTA, 5.95 mM Na_2_ATP, and 2 mM NaN_3_, 5 mM DTT and PIC. *Relaxing-solution* (shortly “*R*” solution; pCa 8.0, ionic strength 170 mM, pH 7.1 at 15°C) contains: 10 mM Imidazole, 2 mM MgCl_2_, 2 mM MgNa_2_ATP, 3 mM EGTA, 110 mM K-propionate (Sigma), 10 mM Caffeine (Sigma), 10 mM Na_2_CrP, 2mM DTT. *Ca^2+^-activating-solution* (ionic strength 170 mM, pCa 4.18, pH 7.1 at 15°C) contains: 10 mM Imidazole, 2 mM MgCl_2_, 2 mM MgNa_2_ATP, 3.1 mM Ca-EGTA (Sigma/Merck), 110 mM K-propionate, 10 mM Caffeine, 10 mM Na_2_CrP, 2mM DTT. The *Standard Ca^2+^-activating-solution* (shortly “*A*” solution; pCa 4.63) has the same composition as the one above, except that Ca-EGTA was at 3 mM. Protease Inhibitor Cocktail (PIC) contains: 1mM AEBSF, 10µM Leupeptin, 10µM Antipain, 1µg/mL Aprotinin and 10µM E64.

Calcium concentration is given as pCa=-log_10_[Ca^2+^]. Solutions with different pCa were obtained by mixing appropriate volumes of *Relaxing-solution* (pCa 8.0) and *Standard Ca^2+^-activating-solution* (pCa 4.63) as indicated below; the free Ca^2+^ concentrations were calculated as previously described in Fabiato and Fabiato, 1979 [4]: pCa 5.24 (95% “*A*”, 5% “*R*”), pCa 5.54 (90% “*A*”, 10% “*R*”), pCa 5.74 (85% “*A*”, 15% “*R*”), pCa 5.80 (83.2% “*A*”, 16.8% “*R*”), pCa 6.12 (70% “*A*”, 30% “*R*”), pCa 6.31 (60% “*A*”, 40% “*R*”).

#### Preparation of ventricular myofibrils

*In vitro* myofibril contractile model, reflecting the coexistence of mutant and wild-type myosin motors in sarcomeric environment, provides a useful framework for investigating mutation-induced alterations in cross-bridge kinetics and for testing hypotheses regarding the effects of HCM-related mutations and myosin modulators at sub-cellular, sarcomeric level.

Myofibrils were isolated either from patient tissue or from three different non-transplanted donor hearts for which no suitable recipient was found and results of experiments pooled together. Donor ventricle myofibrils were considered as control group. Long-term cryopreservation of muscular tissues was previously described [7]. Small pieces of the deep-frozen tissue were generated by slightly crushing the larger biopsies with a mortar and pestle permanently cooled by liquid nitrogen. After quick thawing in *Na-solution* including 20 mM BDM, myocardial fragments were cut in few even smaller pieces, removing connective tissues and fat traces in ice cold *Na-solution* (incl. 20 mM BDM), and then demembranated in cold (∼5°C) *K-solution* containing 1% Triton-X-100 and 20 mM BDM for about 50 minutes. Probes were rinsed three times with cold *K-solution* (+BDM) without Triton (∼50 min) and the third time in absence of BDM. Myofibril-bundles were prepared freshly before the micromechanical experiments by homogenizing the demembranated ventricular pieces equilibrated with cold fresh *K-solution* without BDM for 8-10 s with a blender (Ultra-Turrax T8, IKA Labortechnik, Germany) at 25,000 rpm. The homogenate was further centrifuged (400×g, 10 min, 5°C; Biofuge Primo-R, Thermo-Fisher, Massachusetts, USA) and the pellet re-suspended in fresh *Relaxing-solution* (+PIC, +DTT, pCa 8.0) or subjected to further treatments. The resulting myofibrillar suspension was filtered through a mesh (pore size 21 µm) to remove thick myofibrillar aggregates.

#### Samples treatments

Inotropic drugs administration before organ donation activates PKA via β-adrenergic pathway, while chronic β-blocker therapy reduces cAMP-PKA pathway activation in HCM and end-stage HF patients, diminishing phosphorylation of canonical PKA targets. Therefore, both patient myofibrils (MT-hvMFs) and donor myofibrils (WT-hvMFs) were incubated for 45 min at 20°C with the activated catalytic subunit of protein kinase A (285 U/mL PKA from bovine heart; Sigma-Aldrich, P2645-400U) in *Relaxing-solution* (+DTT, +PIC, pH 7.1) prior functional assessments to minimize the potential effects of different phosphorylation between donor and patient myofibrils. After incubation, myofibrils were rinsed with *Relaxing-solution* (+DTT, +PIC) without PKA. Some myofibrils received the same treatment but without PKA in the *Relaxing-solution* (e.g., in Fig. 4B-F).

Mavacamten (MYK-461 from Cayman Chemical, Cay19216-1), was dissolved in DMSO (dimethyl sulfoxide, Sigma) according to the manufacturer’s instructions. A stock solution of 10 mM mavacamten (mava) was prepared. In experiments, myofibrils were exposed to mavacamten at a final concentration of 0.6 µM (0.001% DMSO concentration) present in both, *Ca^2+^-activating-* and *Relaxing-solutions*; DMSO was present also in control solutions (i.e., without mavacamten) but only in experiments with associated results presented in Fig. 7,8. The procedure was previously detailed [9].

#### Histology and immunostaining

Formalin-fixed paraffin-embedded (FFPE) cardiac tissue samples were sectioned at 2 µm thickness using a microtome (Leica RM 2145). Sections were mounted on glass slides, deparaffinized in xylene and rehydrated through a graded ethanol series.

For general tissue morphology and cross-sectional area (CSA) measurements, sections were stained with hematoxylin and eosin (H&E) following standard protocols (Fig. 2A,B). Cardiomyocyte (CM) size was quantified using a customized version of the CmyoSize ImageJ macro [2] (https://www.sciencedirect.com/science/article/pii/S0940960222000073?via%3Dihub) which enables automated measurement of CM’s CSA from H&E-stained myocardial sections. Primary criteria for inclusion were intact cross-sectional morphology, visible cell boundaries, and a single centrally located nucleus per cell.

The original macro from da Silva et al., 2022 [2] was adapted to optimize segmentation for HCM tissue and to streamline result handling (CmyoSize_TH-2025.ijm). Specifically, the prominence threshold in the “Find Maxima” step was increased (from 5 to 12) to improve cell boundary separation in densely packed HCM myocardium, and morphological smoothing was reduced (erosion/dilation radius adjusted from 10 to 5 pixels) to preserve cell shape fidelity. The area inclusion range was broadened (100 to infinity) to accommodate hypertrophic CMs. Images were analyzed in Fiji (v1.54p) using the Shape Filter and IJBlob plugins. All results obtained with this tool were manually verified. Incorrectly segmented cells were corrected, and additional CMs were manually added to the region of interest (ROI) where necessary. For CSA analysis, a total tissue area of 2.63 mm² was evaluated in donor samples and 11.44 mm² in patient samples. Analyses were based on eight donor and three patient tissue sections (left ventricle).

To assess fibrosis, additional sections were stained with Masson-Goldner (MG) (Fig. 2A,C). All slides were scanned using a high-resolution digital slide scanner (Metafer; MetaSystems Hard& Software GmbH) for subsequent image analysis. Fibrosis and muscle areas were quantified using a custom ImageJ macro (FibroQuant_MG_TH.ijm). The blue/turquoise-stained regions (collagen and connective tissue) were defined as fibrosis, and the red-stained regions as CMs (Fig. 2A). Only these two tissue compartments were included in the analysis; other structures such as fat, vessels, or empty spaces were excluded. The fractional area of fibrosis was calculated relative to the sum of fibrosis and muscle areas. Fibrosis analysis was performed on seven left ventricular tissue sections from donor hearts and three sections from patient hearts. For each donor or patient, a total tissue area of 166,944 µm² (n=8 regions of interest) was analyzed.

For immunofluorescent staining, antigen retrieval was performed in EDTA-containing buffer (pH 9.0) at 99°C in a water bath, followed by permeabilization with 0.5% Triton X-100. Bovine serum albumin (5% BSA) was used for blocking and antibody dilution. Primary antibodies for α-Actinin (ab137346, Abcam; rabbit, polyclonal) and N-cadherin (ABIN1440032, antibodies-online GmbH; goat, polyclonal) were incubated simultaneously. Wheat germ agglutinin (WGA; ab20528, Abcam; conjugated with FITC) was incubated together with secondary antibodies (Alexa 555: A3157, Invitrogen, Thermo Fisher Scientific; and Alexa 680: A32860, Invitrogen, Thermo Fisher Scientific; donkey anti goat) and DAPI (D9542; Sigma). Images were acquired using a Leica TCS SP8 confocal system coupled to an inverted DMi8 microscope.

#### Analysis of sarcomeric proteins

Sarcomeric protein analysis was performed with myofibrillar pellet of adult human ventricular, atrial or skeletal muscle (M. gastrocnemius) samples. All myocyte/muscle samples were demembranated with detergent (1% Triton-X-100).

Coomassie-stained SDS-page gels for detection of sarcomeric proteins ranging from myosin heavy chains (MyHC) (8% SDS gel, with 5% glycerol) [12] to myosin light chains (MLC) (10% or 12% SDS Gel) and for detection of βMyHC and αMyHC isoforms were previously described [7]. For gels, the detection lower limit is 5%. Identity of the sarcomeric proteins and their isoforms was either confirmed by western blotting using primary antibodies against: troponin T (TnT; Abcam Clone TT98, Cambridge, UK), troponin I (TnI; Meridian Clone B2541M, Memphis, Tennessee, USA), tropomyosin (Tm; Sigma Clone TM311, Deisenhofen, Germany), essential light chain (MLC-1a/v; PA5-49205, Thermo Fisher Life Science, Life Technologies GmbH, Darmstadt, Germany), and regulatory light chain (MLC-2a/v; ab48003, Abcam Limited Cambridge, UK) and/or by the size determined using a protein ruler (PageRuler, Thermo-Fischer, Massachusetts, USA) and Colormarker (BioRad Precision, plus Protein Standard for Westernblot, München, Germany), and/or by direct comparison to the known protein isoform type found in human ventricle, atrium and gastrocnemius muscles. For immunoblot analysis, conjugated secondary antibodies and methodological details were previously explained in [13]. Intensity of some protein isoform bands was assessed with LAS4000 software (GE Healthcare, Solingen, Germany) and analyzed with Totallab TL100 software (Nonlinear Dynamics Ltd. Newcastle, UK).

#### Myofibrils micromechanical setup and derived parameters

Myofibrillar homogenate in *Relaxing-solution* (100-300µL) was pipetted in the pre-cooled chamber (15±0.5°C) of the custom-build micromechanical setup and myofibrils allowed to sediment for ∼1 h. Then the chamber was filled with *Relaxing-solution*. A single thin myofibril bundle was mounted between the tips of a piezo-driven (P-841.20, Physik Instrumente, Waldbronn, Germany) stiff needle (AMS5775, Science Products GmbH, Hofheim, Germany) and a nN-sensitive force probe (stiffness 1.2-1.8 nN/nm; SD-ZEIL-10, NanoAndMore GmbH, Wetzlar, Germany) (Fig. 4A). A laser beam (Schäfter+Kirchoff, 51nanoFI, 3B) is reflected by the back side of the force probe onto a motion detector. This architecture allowed to determine the deflection of the cantilever in response to the force exerted by the contracting myofibrils (Fig.4A). Myofibrils were firmly adherent to the tips pre-coated with a 50% (v/v) mixture of flowable silicone rubber (3140 RTV, Dow Corning, Wiesbaden, Germany) with 1% nitrocellulose in amyl-acetate. Myofibrils micromecanical setup was built on an inverted microscope (Olympus IX71) equipped with a CCD camera (Olympus U-CMAD3, Japan). Images with myofibrils (as in Fig. 4A) were taken during the experiment with the Olympus’ objective 60x/0.70 Ph2 through a magnifier lens of 1.6x. The setup has three motorized micromanipulators (MP-285) which drives the stiff needle, force sensor and another sharp tool used to mount and discard the myofibrils.

Before Ca^2+^-activation, sarcomere length (*SL*) of the mounted myofibrils was adjusted to 2.3 µM from their initial slack length (*SL*_0_=1.8-2.1µm); the bundle was then slacked for a defined time to measure the resulting passive force (*F*_pas_). In some experiments, myofibrils were kept in *Relaxing-solution* (pCa 8.0) and stretched and slackened successively to different *SL* up to 2.7µm above their *SL*_0_ and the resulting *F*_pas_ recorded and plotted (as in Fig. S1).

For measurements of the active force (*F*_act_), isometrically held myofibrillar bundle was exposed to laminar solution flows of either *Relaxing-solution* or *Ca^2+^-activating-solution*. Solution change was induced rapidly (10 ms) by the movement of a theta-style micropipette (TGC150-15, Clark Electromedical Instruments, Reading, UK) driven by a piezo-actuator (P289.40, Physik Instrumente, Waldbronn, Germany) as previously described [6, 10, 11]. To calculate the active force (*F*_act_) caused by cycling cross-bridges in sarcomeres, the passive force (*F*_pas_) was subtracted from the total assessed force (*F*_total_): *F*_act_ = *F*_total_ -*F*_pas_ (Fig. 4A). Each mounted myofibrils were minimally Ca^2+^-activated to keep the force run-down as low as possible. Force calibration was done after each analyzed myofibril.

From the force traces of myofibrils (Fig. 4A, 5A, 6A), the following force-related parameters can be derived: *passive force* (*F*_pas_) while myofibrils are in relaxing solution (pCa 8.0), *activation half-time* (*t*_1/2,act_) of Ca^2+^-induced force development while myofibrils are exposed to Ca^2+^-activation solution (either at saturating calcium concentration [Ca^2+^] = pCa 4.18–pCa 4.63 or at submaximal [Ca^2+^] = pCa 5.54), steady-state *isometric force* (*F*_act_) while myofibrils are exposed to solutions with different [Ca^2+^] (between pCa 4.18 and pCa 6.31 for a force-pCa relationship), *rate constant of force redevelopment* (*k*_tr_) while myofibrils are steadily exposed to Ca^2+^-activation solutions (pCa 4.18 or pCa 4.63 or pCa 5.54) after a quick release-restretch maneuver, *relaxation half-time* (*t*_1/2,rel_) of force decay while calcium concentration to which myofibrils are exposed is rapidly (within 10ms) diminished from saturating [Ca^2+^] to low [Ca^2+^], either at pCa 8.0 (to induce full relaxation) or at pCa 6.31 or pCa 5.89 (to induce partial relaxation), and three specific kinetic parameters related to biphasic force decay during myofibrils relaxation. i.e., *rate constant of force decay during first relaxation phase k*_lin_, *duration of the first relaxation phase t*_lin_, and *rate constant of force decay during second relaxation phase k*_rel_.

#### Data analysis and statistics

Activation half-time (*t*_1/2,act_) represents the time required for the force to rise up to half of its maximum upon Ca^2+^-activation. After the quick release-restretch maneuver of the isometrically contracting myofibrils, the trace of force increase was fitted to a mono-exponential function yielding the rate-constant *k*_tr_ [6, 10, 11].

Overall relaxation can be provided using relaxation half-time (*t*_1/2,rel_) defined as the time required for the force to fall to half of its maximum amplitude. During biphasic myofibrillar relaxation, force decay was fitted by a function consisting of a linear and mono-exponential term yielding the kinetic parameters *k*_lin_, *t*_lin_, and *k*_rel_ [6, 10, 11]. Relaxation half-time is in the following relationship with the other three relaxation kinetic parameters: *t*_1/2,rel_ = *t*_lin_+ln(2)/(*k*_rel_+ln(1-*k*_lin_·*t*_lin_)/*k*_rel_).

Concentration of the free calcium ions [Ca^2+^] was given as pCa=-log_10_[Ca^2+^]. Force unit is given in kPa=nN/µm^2^. In the force-pCa relationship (Fig. 4C), force is normalized to *F*_act,max_ (at pCa 4.18). Fig. 4C shows the fit curves of the pooled normalized force values with the sigmoidal Hill-equation *F*_act_(pCa) = 1 /(1 + 10^(*n*_H_·(pCa_50_-pCa))). Force-pCa relationships resulting from each measured myofibrillar bundle was fit to the Hill-equation and the mean values of the resulting parameters pCa_50_ (i.e., [Ca^2+^] at half of maximum generated force indicating Ca^2+^-sensitivity of isometric force) and *n*_H_ (i.e., the Hill coefficient indicating the steepness of the force-pCa relation) are shown either in the insets of Fig. 4C as mean±SD or as mean±SEM in Table S1.

If not stated otherwise, steady-state and kinetic parameters were shown as mean±SD (SD = Standard Deviation), while in Table S1 and in Fig. 6B (near the top of each bar) as mean±SEM (SEM = Standard Error of the Mean). Mean values were compared using unpaired Student *t*-test if not otherwise stated; considered levels of significance: *p*<0.05 (∗), *p*<0.01 (∗∗), or *p*<0.001 (∗∗∗).

In Figs. 7,8, values in percent (%) represents how much the respective parameter was above (+) or below (-) the corresponding mean value of the left side bar (taken as reference) indicated by the horizontal line.

#### Computer programs

GraphPad Prism program was used for statistical analysis, plotting and fitting the data (GraphPad Software, LLC, version 9.3.1, 471, 2021; https://www.graphpad.com/). To retrieve the elements of myosin secondary structure from Protein Data Banks published by Auguin et al., 2024 (Auguin et al., 2024), the UCSF ChimeraX software (version 1.10.1, 2025-07-24; https://www.rbvi.ucsf.edu/chimerax) was used.

## ABBREVIATIONS

ACE: **a**ngiotensin-**c**onverting **e**nzyme
BDM: 2,3-**b**utanedione **m**onoxime
[Ca^2+^]: **c**alcium **c**oncentration(s)
CM(s): **c**ardiomyocyte(s)
CSA: **c**ross **s**ection **a**rea
DTT: **d**ithiothreitol
EF: **e**jection **f**raction
HCM: **h**ypertrophic **c**ardiomyopathy
HOCM: **h**ypertrophic **o**bstructive **c**ardiomyopathy
hvMFs: **h**uman **v**entricular **m**yofibrils
IHM: **i**nteractive **h**ead **m**otive
LV: **l**eft **v**entricle, **l**eft **v**entricular
LVOT: **l**eft **v**entricular **o**utflow **t**ract
Mava: **m**avacamten
MLC-1/2: **m**yosin **l**ight **c**hain 1 (**e**ssential **l**ight **c**hain, ELC) / **m**yosin **l**ight **c**hain 2 (**r**egulatory **l**ight **c**hain, RLC)
MyBPC: **m**yosin **b**inding **p**rotein C
MyHC: **m**yosin **h**eavy **c**hain
MT: **m**utant
NYHA: **N**ew **Y**ork **H**eart **A**ssociation
PDB: **p**rotein **d**ata **b**ank
P_i_: **i**norganic **p**hosphate
PIC: **p**rotease **i**nhibitor **c**ocktail
PKA: **p**rotein **k**inase A
SL: **s**arcomere **l**ength
TnI, TnT: **t**roponin I, **t**roponin T
Tm: **t**ropomyosin
WGA: **w**heat **g**erm **a**gglutinin
WT: **w**ild-**t**ype

